# Are populations like a circuit? Comparing isolation by resistance to a new coalescent-based method

**DOI:** 10.1101/451328

**Authors:** Erik Lundgren, Peter L. Ralph

## Abstract

A number of methods commonly used in landscape genetics use an analogy to electrical resistance on a network to describe and fit barriers to movement across the landscape using genetic distance data. These are motivated by a mathematical equivalence between electrical resistance between two nodes of a network and the “commute time”, which is the mean time for a random walk on that network to leave one node, visit the other, and return. However, genetic data are more accurately modeled by a different quantity, the coalescence time. Here, we describe the differences between resistance distance and coalescence time, and explore the consequences for inference. We implement a Bayesian method to infer effective movement rates and population sizes under both these models, and find that inference using commute times can produce misleading results in the presence of biased gene flow. We then use forwards-time simulation with continuous geography to demonstrate that coalescence-based inference remains more accurate than resistance-based methods on realistic data, but difficulties highlight the need for methods that explicitly model continuous, heterogeneous geography.

## Introduction

Genetic relatedness is determined by past gene flow, a product of individual or gamete movements across geographic space. Genomes therefore retain the traces of this movement, and can contribute to the inference of how a species moves across a landscape, which is important for understanding how diseases spread, how species adapt, and how to retain genetic diversity in threatened species. This is a potentially important source of information, as direct observation can be difficult (especially of long-distance movement, [Cayuela et al., 2018, Levin et al., 2003]), or even impossible, if some of the populations in question no longer exist.

The idea of “resistance distance” is an important tool in the landscape genomics toolbox. Introduced by McRae [2006], it makes use of a mathematical equivalence between random walks and electrical resistance [Nash-Williams, 1959]: the expected time for a continuous-time Markov chain that starts at node *x* to first hit node *y* and then return to *x* (the “commute time”) is equal to the effective resistance between *x* and *y* in an electrical network whose conductances are given by the movement rates of the Markov chain [Levin et al., 2008, Doyle and Snell, 2006]. This measure therefore averages over all possible paths through the network between the two. Suppose we assign local conductances based on the values of some landscape variable, and compute the correlation of genetic distances between samples from different parts of the landscape with the effective resistances between the samples’ locations. If this results in a significant positive correlation, then this is taken as good evidence that the landscape variable is a good indicator of where gene flow occurs [McRae and Beier, 2007, Cushman et al., 2006]. For instance, if the conductance across a grid cell in a discretization of the landscape is higher in flatter areas, and this produces effective resistance values that are significantly correlated with genetic distances, then one might conclude that the species in question tends to disperse more readily through flatter areas, perhaps along river valleys. More recent methods [Petkova et al., 2016, Hanks and Hooten, 2013] seek to build *de novo* a map of conductance values that produces resistances most strongly correlated with observed genetic distance, and then interpret regions of low conductance in the resulting map as barriers to gene flow.

As motivated by McRae [2006], commute time (i.e., resistance distance) is a computationally tractable approximation to the more accurate random walk-based model of genetic distance, the *coalescence time*. Genetic distances between two genomes sampled from the landscape do derive from an average across a large number of paths between those points – the lineages along which each segment of genetic material has been inherited from their most recent common ancestor. Thanks to recombination, there are a large number of such paths, and genetic distance averages over these. The more recent these common ancestors tend to be, the smaller the genetic distance is. It is reasonable in some situations to model these lineages as random walks across the landscape; however, the model of genetic distance that one is thus led to is *coalescence time* of the random walks, rather than commute time, the quantity that corresponds to resistance. This naturally raises the question: Are methods that depend on effective resistances being misled by model misspecification? If so, would using coalescence time do better?

McRae [2006] showed that coalescence time and resistance distance are equal (up to shift and scaling) in isotropic landscapes such as a ring [Matsen and Wakeley, 2006], and found that the two were highly correlated in several test cases. However, all test cases had symmetric migration (the same rate of gene flow in both directions along each edge), which is likely not the case in many situations, e.g., in rivers [Morrissey and de Kerckhove, 2009, Sundqvist et al., 2016, Hanks, 2017], with source-sink dynamics [Dias, 1996], or after population expansions [With, 2002].

In this paper, we contrast coalescence time and resistance distance, and develop a method to infer movement rates on a discrete landscape using genetic distances. As we will see, although the two methods are conceptually similar, resistance-based inference may infer movement rates uncorrelated with the truth if data are derived from a coalescent process, especially in the presence of gene flow asymmetry. We also explore the question of identifiability. Resistance-based methods are often used to predict distances based on a few layers of geographical data (land cover, slope, etcetera). Some methods find the combination of landscape layers to best explain genetic distance, which allows very fine geographic resolution [Shaffer et al., 2017]. Other methods such as EEMS [“Estimating Effective Migration Surfaces”, Petkova et al., 2016] infer conductances without such prior information; however, the geographical resolution is much coarser. (A recently-released method that is based on EEMS but uses haplotype length data and an underlying model of coalescence time [Al-Asadi et al., 2018] is promising, but still produces similarly coarse maps.) One reason for this difference in resolution is that EEMS uses more computationally intensive Bayesian methods. However, we show that this coarse resolution reflects a more general problem, both because of unmodeled “process noise” involved in discretization of geographical populations, and the mathematical structure of the underlying problem.

## Methods

The *commute time* of a Markov chain between locations *i* and *j* is the expected time for a particle that moves according to the rules of the chain, started at *i*, to first reach *j* and then return. We will denote this by *R*_*ij*_, to emphasize that it is also the effective resistance between *i* and *j*. The *coalescence time* is the expected time for two particles started at *i* and *j* to *coalesce*, which happens at a given rate whenever they are in the same location. We will denote this by *C*_*ij*_. Why should these be related to genetic distance? We first explain this connection and discuss computation of both quantities, and finally develop a method that infers movement rates of the Markov chain (which are rates of gene flow) from observations of coalescence time. We apply the method to both realistic simulations in continuous space and to an empirical dataset. Although we have tried to highlight the intuitive aspects of the initial technical discussion, readers who are interested in practical differences more than the mathematical arguments should be able to skip to “Test cases and simulations” without loss of continuity.

### Genetic distance and coalescence time

There are a number of ways to calculate measures of dissimilarity using genetic data. In this paper we use mean pairwise divergence, which for two genomes is calculated as the proportion of sites in the genome at which the two genomes differ. It is more common in the literature to use *F*_*ST*_ */*(1-*F*_*ST*_), but the two are highly correlated for good theoretical reasons [Slatkin, 1991, Rousset, 1997], especially for SNP data. (However, note that *F*_*ST*_ depends on the sampling scheme, while divergence does not.) The two genomes differ at a site only if there has been a mutation in some ancestor on the lineages leading from the two genomes back to their most recent common ancestor at that site. Therefore, under an infinite-sites model with average mutation rate *µ* per site and per meiosis, pairwise divergence divided by *µ* is an unbiased estimate of the mean time to most recent common ancestor, averaged across the genome [Hudson, 2007, Ralph, 2015]. It is natural to model this process by following the two lineages back through time (and across space) until they find their most recent common ancestor, i.e., until they coalesce. Suppose we model the geographic distribution of the species as a collection of randomly mating populations that exchange occasional migrants, thus discretizing geographic space. It is common to assume that the motion of these lineages back through time forms a Markov process, the *structured coalescent* [Wakeley, 2005]. (This assumption requires the effects of natural selection on each segment of the genome to be generally weak.) In this framework, each lineage performs a random walk across the populations, with movement probabilities depending on the flux of individuals between populations. This provides the link between genetic distance and random walks.

For data, we are given genotypes of individuals and their geographic coordinates; then, we divide these individuals into groups, and compute the genetic distance between group *i* and group *j* as the mean divergence between a pair of genomes, chosen one from each group. We denote the resulting genetic distance *D*_*ij*_; since divergence is symmetric, *D*_*ij*_ = *D*_*ji*_. We compute *D*_*ii*_ by sampling without replacement from group *i*, so that generally *D*_*ii*_ > 0: the diagonal elements measure local diversity. As discussed below, “resistance distance” methods find commute times so that the mean commute time between *i* and *j* matches *D*_*ij*_ – but the commute time of a location to itself is zero, so a term is added to account for local diversity. We do this in the same way as Petkova et al. [2016], by introducing the *local diversity* parameters *q*_*i*_, and using the model

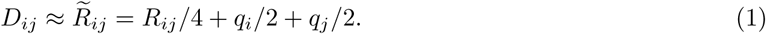

Concretely, this means finding movement rates of a Markov chain whose matrix of commute times (*R*) yields a good approximation to the matrix of genetic distances, after adding local diversity parameters (*q*, which are also free parameters). The factor of four is necessary to make commute time work on the same time scale as coalescence time: the commute time covers the path from *i* to *j* twice (once in both directions), while the coalescence time covers the distance only once, and in half the time, because two particles are moving. For a conceptual picture of the differences between the coalescence time and commute time, see Figure 1.

**Figure 1:**
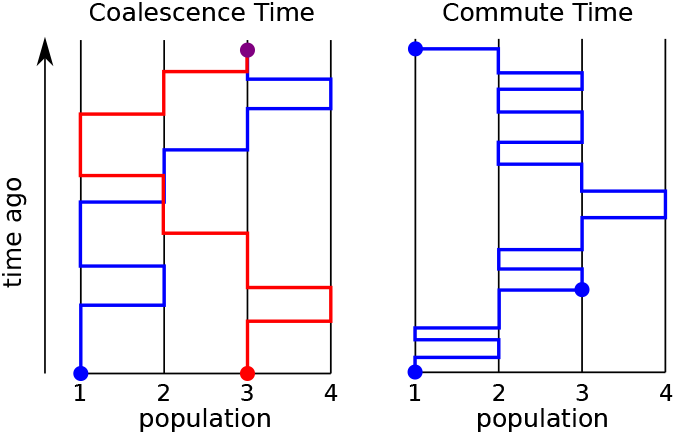
Illustration of the conceptual differences between coalescence and commute time between states 1 and 3 for a continuous time Markov chain with four linearly connected states. Coalescence time (left) is the time for two independently moving particles to meet and coalesce (it is possible for them to be in the same location without coalescing). Commute time is time for a single particle starting in one state to travel to another state and then back to the original state. Note how the time axis is scaled to be faster in the figure showing commute time than the one for coalescence time in order for them to fit in the same space.

A simple situation when the two clearly differ is in the presence of biased gene flow, as for instance between a source and a sink population: coalescence only requires that lineages move in *one* direction between populations, while a commute requires movement in both.

### Hitting times of Markov chains

Now, we explain how to compute the relevant quantities – mean commute and coalescence times – from a Markov chain model of lineage movement. The Markov chain is defined by its *generator matrix*, denoted *G*, for which *G*_*xy*_ gives the “jump rate” from *x* to *y*. The jump rate *G*_*xy*_ is the probability the chain jumps to position *y* when at position *x* per unit of time. (More precisely, if the Markov chain is at location *X*_*t*_ at time *t*, then for *x≠y*, ℙ{*X*_*t*+ϵ_= *y* | *X*_*t*_ = *x*} = ϵ*G*_*xy*_ + *O*(ϵ^2^). For mathematical convenience, the diagonal of this matrix is completed so that rows sum to zero: *G*_*xx*_ = *-*Σ_*y*≠*x*_ *G*_*xy*_. We will need to find the *hitting times* of the chain – i.e., for each pair of states *x* and *y*, the mean time until the chain first hits *y* after being started at *x*. We denote this quantity *H*_*xy*_. Throughout, we assume that all hitting times are finite, which implies the chain is connected and irreducible. The Markov property lets us write a system of linear equations that can be solved for *H*, as follows (see, e.g., Kemeny and Snell [1983]): Suppose that the chain begins at *x ≠ z*, and is at location *y ≠ z* after *dt* units of time, without having visited *z*. Then the total time to hit *z* is equal to the elapsed time *dt* plus the remaining time to hit *z* from *y*. Taking the average over possible paths, this says that *H*_*xz*_ *≈ dt* + Σ_*y* ℙ_{*X*_*dt*_ = *y* | *X*_0_ = *x*}*H*_*yz*_. Replacing _ℙ_{*X*_*dt*_ = *y* | *X*_0_ = *x*} with *dtG*_*xy*_ and rearranging gives the system of equations:

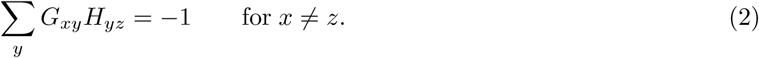

In other words, if *G*_*-z*_ is the matrix with the *z*^th^ row and column removed, and **1** is the vector composed of all ones, then the *z*^th^ column of *H*, except for *H*_*zz*_, can be computed as *-*(*G*_*-z*_)^*-*1^**1**. This, along with *H*_*zz*_ = 0, allows computation of *H* from the movement rates *G* [Kemeny and Snell, 1983].

Suppose instead we are given the hitting times, *H*, and wish to find the movement rates, *G*. This is not the situation we are in – we have either commute or coalescence times – but it is related. The Random Target Lemma [Aldous and Fill, 2002] tells us that the stationary distribution of the Markov chain, denoted *π*, can be recovered from the hitting times by solving π = *H*^*-*1^**1***/***1**^*T*^ *H*^*-*1^**1**. As shown in Appendix B, equation (2) can be rewritten in matrix form as

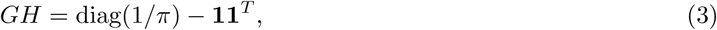

which implies that *G* can be computed directly from *H* as

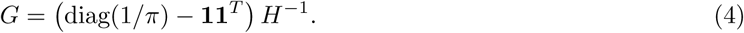

(We prove that *H* is invertible in Appendix B.) Therefore, the pairwise mean hitting times, *H*, uniquely determine the jump rates, *G*.

The *commute time* between *x* and *y* is the mean time a chain started at *x* takes to first hit *y* and then to return to *x*. We will write this as

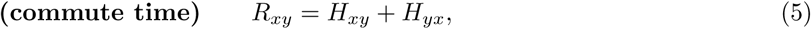

or in matrix notation, *R* = *H* + *H*^*T*^. Note that the commute times are symmetric, even though the hitting times may not be.

Although commute times are uniquely determined by movement rates, the reverse is not true – we can use equations (3) and (5) to see how to modify movement rates without changing commute times. A small concrete example of this is given in Appendix A. More generally, commute times only depend on the symmetric part of *H* – given a skew-symmetric matrix *Z* (so that *Z* + *Z*^*T*^ = 0), any Markov chain that has hitting times given by *H*_*ϵ*_ = *H* + *ϵZ* for some number *ϵ* will have the same commute times. The resulting hitting times may not be valid – if *G*_*ϵ*_ is the matrix constructed by applying equation (4) to *H*_*ϵ*_, then it is not guaranteed that the offdiagonal elements of *G*_*ϵ*_ are all nonnegative, as required. However, since *G* and *H* are continuous functions of each other, if all offdiagonal elements of *G* are strictly positive, then there exists a positive *ϵ*_0_ such that all *G*_*ϵ*_ for *ϵ* < *ϵ*_0_ do define valid Markov chains, all with the same commute times.

The *coalescence times* are defined using *two* copies of the same chain, as the mean time until coalescence, if the chains coalesce at rate *γ* when they are in the same place. Concretely, suppose that *X* and *Y* are independent Markov chains both moving with movement rates given by *G*, that coalesce at rate *γ*_*x*_ when *X* and *Y* are both at *x*. Define *τ* to be this coalescence time, so that ℙ {*τ*≤*t* + *ϵ τ* > *t, X*_*t*_ = *Y*_*t*_ = *x*}= *ϵγ*_*x*_ + *O*(*ϵ*^2^), and ℙ{*τ* ≤*t* + *ϵ*|*τ* > *t, X*_*t*_ ≠ *Y*_*t*_} = *O*(*ϵ*^2^). Then, the (mean) coalescence time is defined to be *C*_*xy*_ = 𝔼 [*τ X*_0_ = *x, Y*_0_ = *y*]. An argument similar to the one behind equation (2) shows that the coalescence times satisfy the following system of equations:

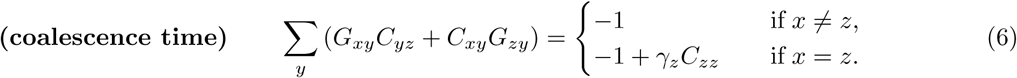

Similar recursions are common in the literature, going back at least to Hill [1972] (see also Whitlock and Barton [1997]). In matrix notation, this is

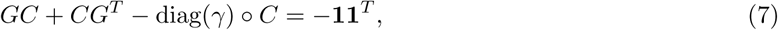

where diag(*γ*) is the matrix with the vector *γ* on the diagonal and zeros elsewhere, and ◦ is the componentwise product. In practice, we solve this by working with the product Markov chain (*X*_*t*_, *Y*_*t*_), whose generator matrix is *G* ⊗ *I* + *I* ⊗ *G*, where *I* is the identity matrix and ⊗ is the Kronecker product.

Equation (7) is linear in *G* and *γ*, and so after rearrangement can be solved with standard linear algebra (and is related to the Sylvester equation [Bhatia and Rosenthal, 1997]). However, the solution is again not necessarily unique: suppose that we have a matrix *Z* such that *ZC* is skew-symmetric. Then (*G* + *Z*)*C* + *C*(*G* + *Z*)^*T*^ = *GC* + *CG*^*T*^, and so a Markov chain with generator matrix *G* + *Z* has the same coalescence times as the original chain. Furthermore, for *G* + *Z* to be a valid generator of a Markov chain, the rows of *Z* must sum to zero, and all offdiagonal entries of *G* + *Z* must be nonnegative. As before, if all entries of *G* are nonzero, it is always possible to find sufficiently small *ϵ* so that *G* + *ϵZ* remains the generator of a valid Markov chain. The coalescence rates, *γ*, provide additional degrees of freedom which may render solutions nonunique in a broader range of situations.

#### Constraints and uniqueness

We have seen that to find the movement rates of a continuous-time Markov chain given the coalescence or commute times we must solve either equation (7) or equations (4) and (5) for *G*, and that these do not have unique solutions. The situation is worse when one needs to infer coalescence rates (*γ*) as well. However, usually in applications the locations lie across geographical space, so that many of the entries of *G* can be assumed to be zero. This reduces the number of unknowns to solve for, and so can render the solution unique. We can get an idea of this by simply counting equations and unknowns. For concreteness, suppose that the spatial locations are arranged in a square grid of *n* locations, so that each is connected to four others. (Boundary locations will have fewer, but we omit this detail.) Since movement rates in each direction can be different, there are 4*n* free parameters, each corresponding to an off-diagonal entry of *G*. Coalescence rates provide another *n* parameters.

Since coalescence times are symmetric, equation (7) provides *n*(*n* + 1)*/*2 informative equations. This is larger than 4*n* + *n*, the number of parameters, for a grid with at least nine nodes (i.e., 3×3), so we would expect the system of equations to usually have a unique solution for grids larger than this, although some special cases will still have free parameters. The same calculation holds for commute times except that there are *n* local diversity parameters instead of *n* coalescence parameters.

This suggests that nonuniqueness of solutions may not be a problem in practice, as long as geography is discretized into sufficiently many regions and direct, long-distance migration is disallowed (i.e., parents and offspring live in the same or neighboring regions). However, finer geographic resolutions may present problems of nonidentifiability, similar to the problem of multiple linear regression with many collinear variables. The problem of inferring gene flow rates on fine geographic maps may become *ill-conditioned*, i.e., arbitrarily small variations in the data (even numerical instability) may produce large differences in the inferred “best” rates. Information about the landscape contained in higher-order modes (e.g., finer resolution changes in hitting times) is obscured by noise. (For the interested technical reader, this could occur as the spatial resolution increases because the matrix *G* converges to a second-order elliptic differential operator [Stroock and Varadhan, 1997]; such operators are deformations of the Laplacian [Feller, 1957], which has rapidly decaying eigenvalues [McKean and Singer, 1967, Kuttler and Sigillito, 1984].) This implies a fundamental limit to the geographical resolution of inferred maps. For more discussion of related problems, see Epstein and Schotland [2008], Myers et al. [2008], or Terhorst and Song [2015].

One strategy for circumventing ill-conditioning is to impose additional constraints – for instance, constraining all population sizes to be the same (something we do below), or allowing only two distinct migration rates: one rate across a barrier, and another across “non-barriers”. Such constraints amount to a form of regularization, since they look for solutions to the original problem with certain desirable properties (e.g., most of the values are the same).

#### When are coalescence and commute times equal?

If in a given biological situation commute and coalescence times are equal – i.e., if given *G* and *γ*, there was a *q* that made 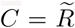– then inference with either model would be equivalent. This was demonstrated in some situations by McRae [2006]. We have already shown that this is not the case in general, but it turns out that it *is* true under some restrictive assumptions commonly found in abstract population genetics models. We show in Appendix C that if hitting times are symmetric (*H* = *H*^*T*^), then this does occur, and so the two methods are equivalent for symmetric island models (where the populations are arranged on a ring and migration rates only depend on the distance between them). Hitting times are not symmetric for a square grid with uniform migration (the mean time to hit the center from a corner is less than the reverse), but the square grid is quite close to the torus, which does have symmetric hitting times.

### Bayesian inference of movement rates

Mean coalescence times estimated from real data are subject to a number of sources of noise (which we discuss in more detail later). Here, we use exact solutions of the equations above to develop a Bayesian inference method that accounts for noise in the data and estimates uncertainty in the resulting estimates. To do this, we model genetic distances as equal to coalescence times plus noise: *D*_*ij*_ = *C*_*ij*_ + *ϵ*_*ij*_, where *ϵ*_*ij*_ are independent and Gaussian distributed with mean **0** and variance 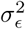, except that *ϵ*_*ij*_ = *ϵ*_*ji*_. (We assume that coalescence times are measured to good accuracy – i.e., *C*_*ij*_ is large compared to *σ*_*ϵ*_ – so that this model does not predict negative times.) Given *G* and *γ* we can solve equation (7) to find the corresponding coalescence times – which we call 𝒞 (*G, γ*) – so our model is that *D* has a Gaussian distribution with mean 𝒞 (*G, γ*). (This Gaussian stands in for the sampling noise of coalescence times about their mean.) The maximum likelihood estimate of *G* could be found directly; however, this in practice will quite likely contain negative movement rates, and does not allow us to impose constraints (such as only allowing a subset of movement rates to be nonzero).

Therefore, we use Bayesian inference, placing independent exponential priors on the parameters – the movement rates, *G*_*ij*_, that are not constrained to be zero and the coalescence rates, *γ*. Concretely, suppose that the spatial arrangement of populations is given by a graph (i.e., a discrete collection of locations represented by “nodes”, between which movement is possible only between those connected by “edges”). Only movement rates in *G* that correspond to edges in the graph are allowed to be nonzero. Suppose there are *m*_*G*_ edges in the graph, that each of the *n* populations has its own coalescence rate, and write *i* ∼*j* to mean that *i* and *j* are adjacent in the graph. Then, the resulting log-posterior density is

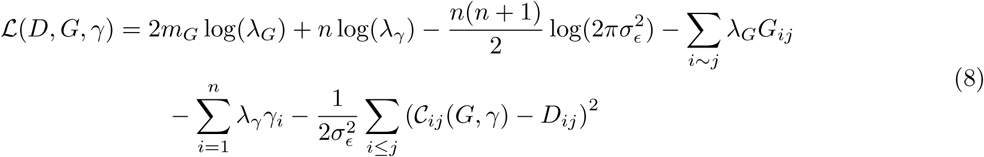

Here 1*/λ*_*γ*_ and 1*/λ*_*G*_ are the prior means of *γ*_*i*_ and *G*_*ij*_, respectively. Recall that both *D* and (*G, γ*) are full matrices of pairwise distances between discrete locations (in the same order), and so the term (𝒞_*ij*_(*G, γ*) − *D*_*ij*_)^2^ measures the deviation of the observed genetic distances from those predicted under the coalescent model on the graph.

Inference with resistance distance is done in exactly the same way, except that *q* replaces *γ*, and instead of 𝒞(*G, γ*) we use 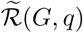, which is the matrix 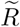 computed from *G* and *q* by putting the solution to equation (2) through equations (5) and (1).

It is not required in this approach to have samples from every spatial location, allowing inference on incompletely sampled spatial discretizations. (However, more complete sampling will produce more accurate results.) To carry out inference in this situation, we simply use Equation 8, but only sum over observed entries of *D*_*ij*_.

#### MCMC methods

To sample from the posterior distribution of equation (8), we used R [R Core Team, 2018] to implement a standard Metropolis-Hastings algorithm with Gaussian proposals reflected into the positive quadrant [Brooks et al., 2011]. Starting locations were chosen by sampling from the prior distribution. Before the usual “burn-in” phase, we included a period of “pre-burn-in” which used the same MCMC procedure with a larger *σ*_*ϵ*_, to allow the chain to more quickly converge on the high-posterior portion of parameter space. Typically, we ran MCMC for 10^6^ iterations of pre-burn-in and 3×10^6^ iterations of burnin, followed by 4×10^6^ iterations which were used to estimate posterior distributions, which took around 10 hours on a single core of a modern computer with 16 demes. For small graphs fewer iterations were necessary. We used standard methods to assess convergence and mixing.

### Test cases and simulations

We compared inference under coalescence time and resistance distance models using both data from the model and from more realistic simulations. For the first category, we designed a number of graphs to test particular aspects of inference, with migration rates specified on each directed edge (plotted below using igraph [Csardi and Nepusz, 2006]). Given the edge weights (*G*) and the coalescence rates (*γ*) of a graph, we produced data by calculating exact coalescence times (𝒞 (*G, γ*)), to whose entries we added independent Gaussian noise while preserving symmetry. We scaled the noise to the mean value of the times themselves, so that a “noise level” of *α* refers to data in which the standard deviation of the noise was set to *α* multiplied by the average coalescence time across all pairs of populations. Particular parameters and levels of noise are given in the Results. These simulations are expected to be nearly equivalent to neutral, discrete population-based simulations (either forwards-time or coalescent), except that noise terms about the theoretical mean should be somewhat correlated.

#### Forwards-time simulations

To produce realistic data, we implemented forwards-time simulations using SLiM v3.1 [Haller and Messer, 2018, **?**], with individuals living across continuous, two-dimensional geography (sometimes with barriers), from which we recorded genomic data. The basic simulation, which we modified to produce several other situations, is as follows. Each simulation had around 6400 diploid, hermaphroditic individuals, living across two-dimensional geography. Mates were selected from nearby individuals using a Gaussian kernel with standard deviation *σ*_*d*_, truncated at 3*σ*_*d*_. Simulations were done on either a square region with width equal to 80*σ*_*d*_, or a 5 : 3 rectangular region with the same area. Offspring are dispersed away from the mother using the same kernel, reflected from the boundaries of the habitable area. To produce a realistic model and prevent excessive population clumpiness [Felsenstein, 1975], we model local competition instead of a globally constant population size. To do this, we use the same truncated Gaussian kernel to define an “interaction strength” between each pair of nearby individuals, and compute the probability of survival of each individual to the next time step as min(0.9, 2*/*(1 + 5*x/*12*π*)), where *x* is the sum of the interaction strengths of that individual with all other individuals within distance 3*σ*_*d*_. (The value *x* is an estimate of local population density.) This reduces fitness for individuals in denser areas, and produces an equilibrium population density of close to one individual per 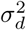 units of area. The probability of survival for individuals within 10*σ*_*d*_ of the edge of the habitat was multiplied by a linear factor that scaled from 0 to 1 with distance from the edge. Patchiness of the resulting simulations was similar to what is seen in Figure 6.

**Figure 2:**
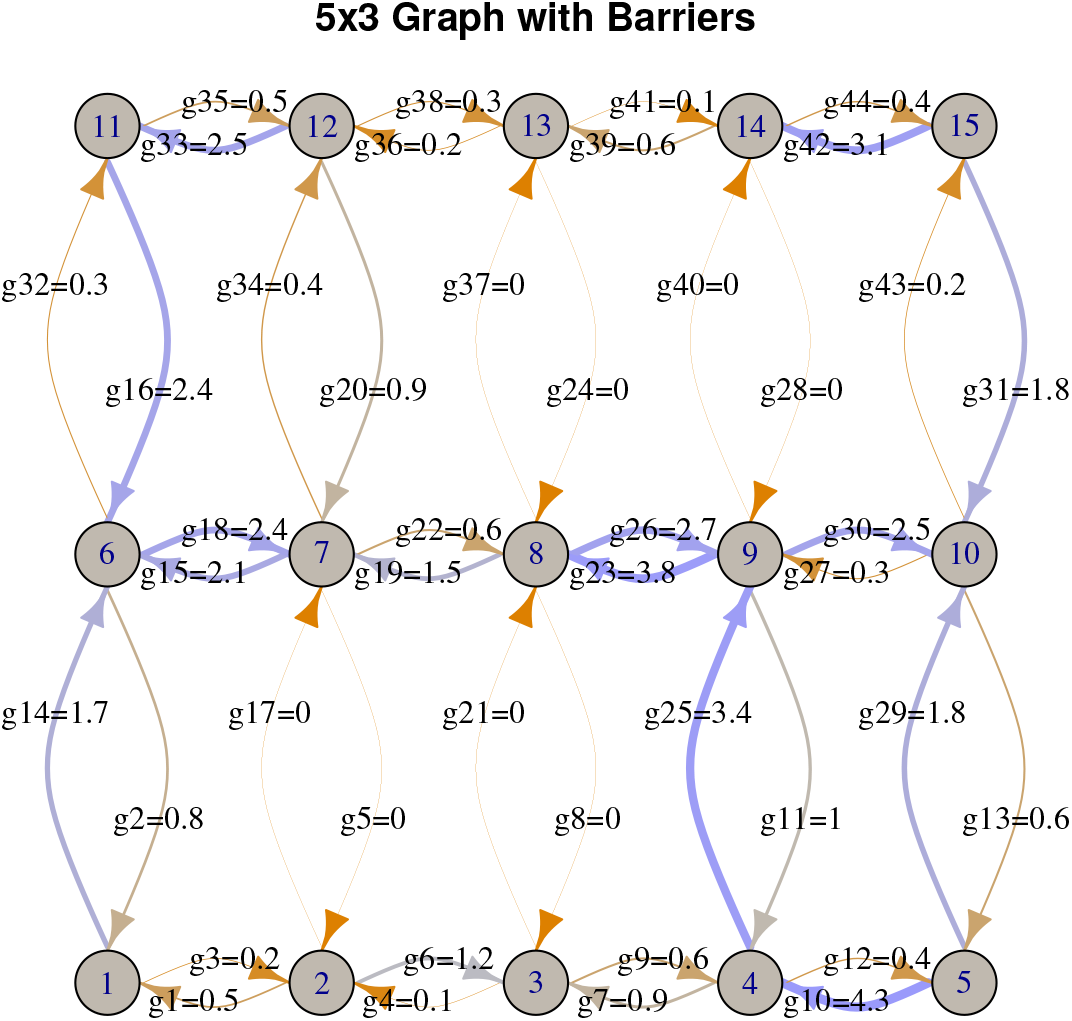
The grid structure and parameter labels for the 5×3 grid with two barriers: movement rates between {2, 3} and {7, 8} are zero, as are those between {8, 9} and {13, 14}. Values for the non-zero movement rates were chosen by rounding independent draws of an exponential random variable with mean 1 up to the nearest tenth. Thickness and color of arrows reflect movement rates: darker red arrows have smaller rates and darker blue arrows have larger rates.

**Figure 3:**
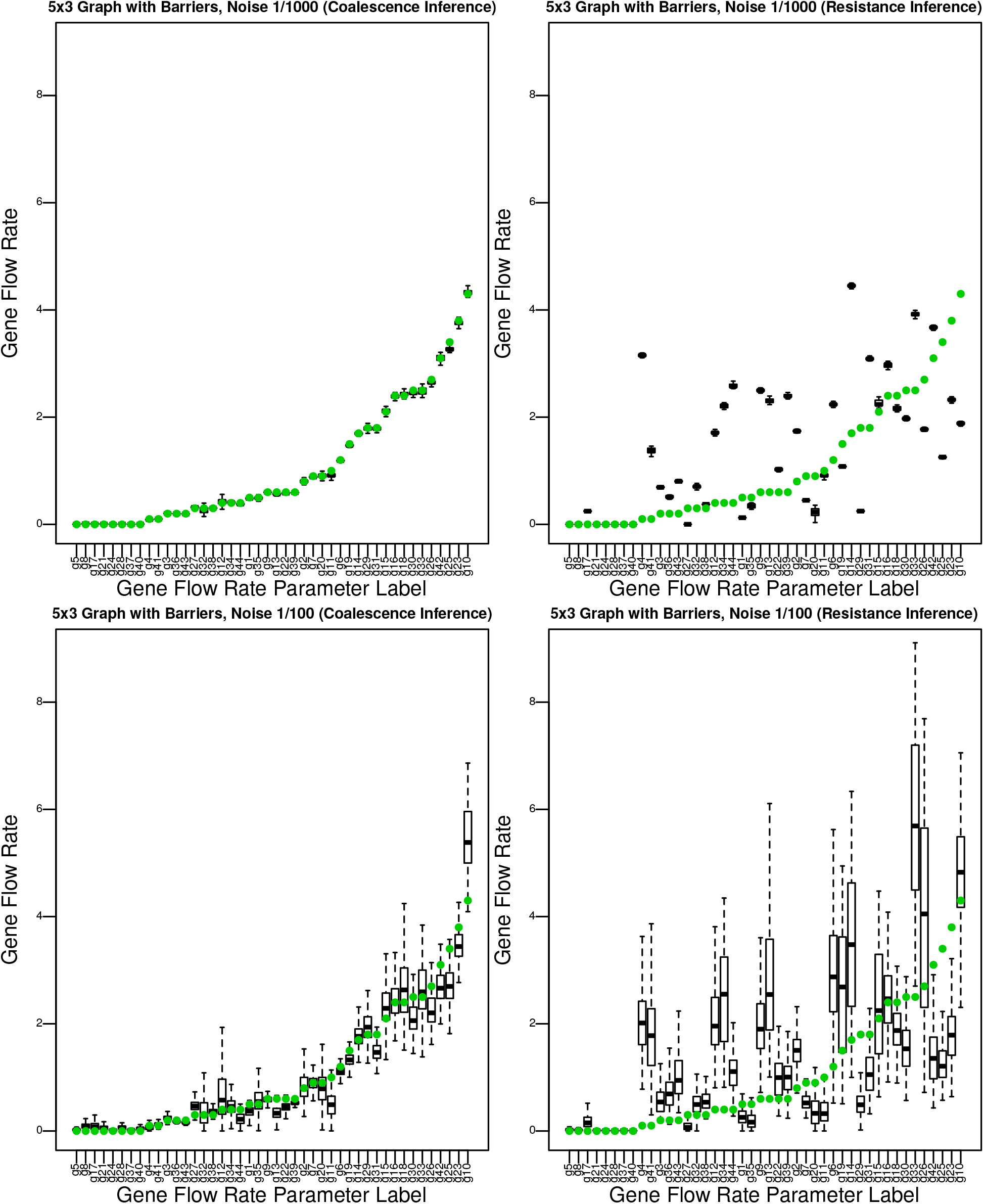
Coalescence time inference is much more accurate than resistance inference: posterior distributions for gene flow rates in the 5×3 graph with barriers, compared to true values (in green), using **(left)** coalescence time and **(right)** resistance distance inference, with **(top)** low noise and **(bottom)** high noise, See Figure 2 to match parameter labels to positions on the graph.

**Figure 4:**
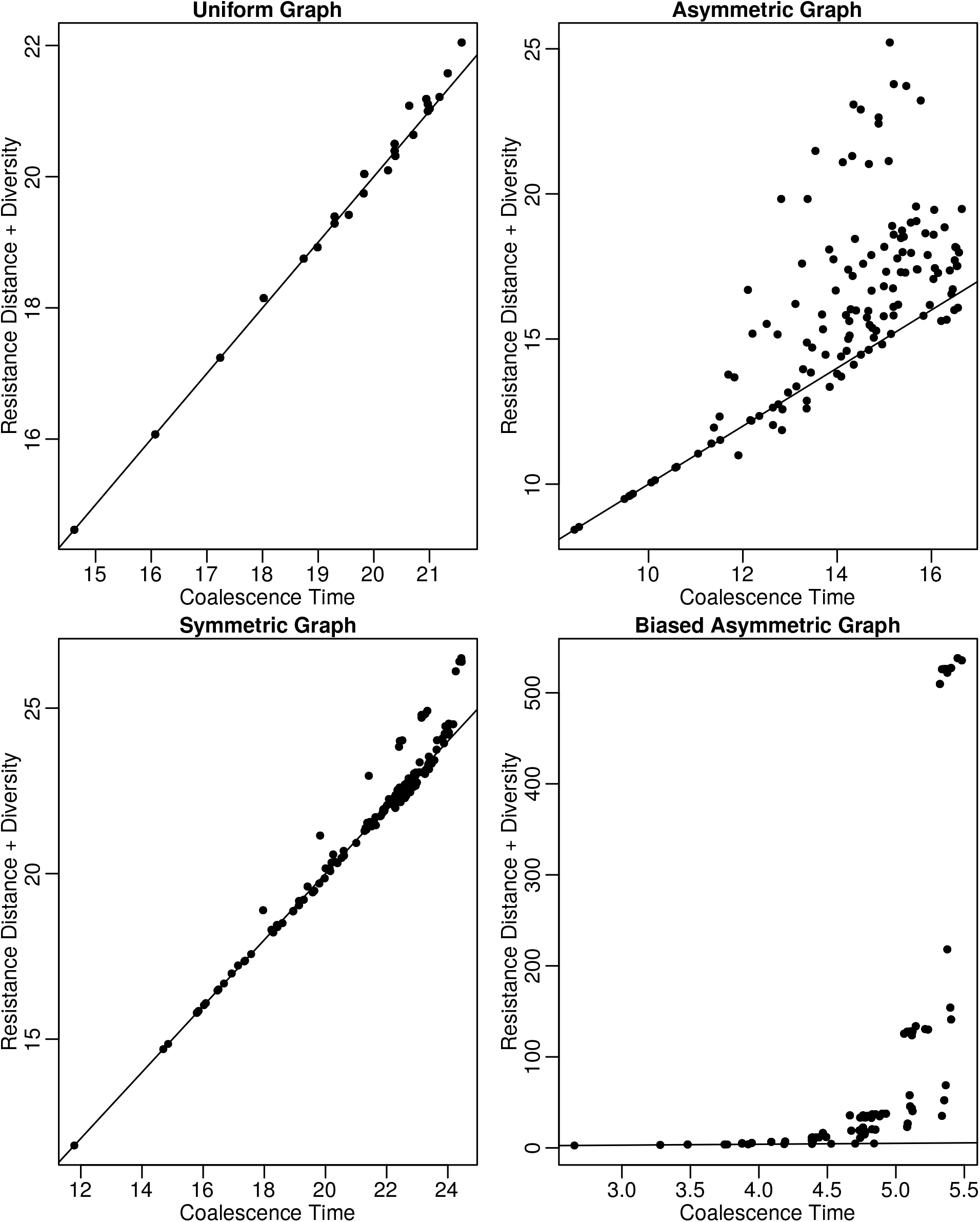
Resistance distances 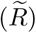 compared to coalescence times (*C*) on the 4×4 graph, with **(top left)** uniform movement rates, **(bottom left)** random, symmetric rates, **(top right)** random, asymmetric rates, and **(bottom right)** rates with a small diagonal bias (see text). The line shows *y* = *x*. See Figure S7 for specific values and graph structure.

**Figure 5:**
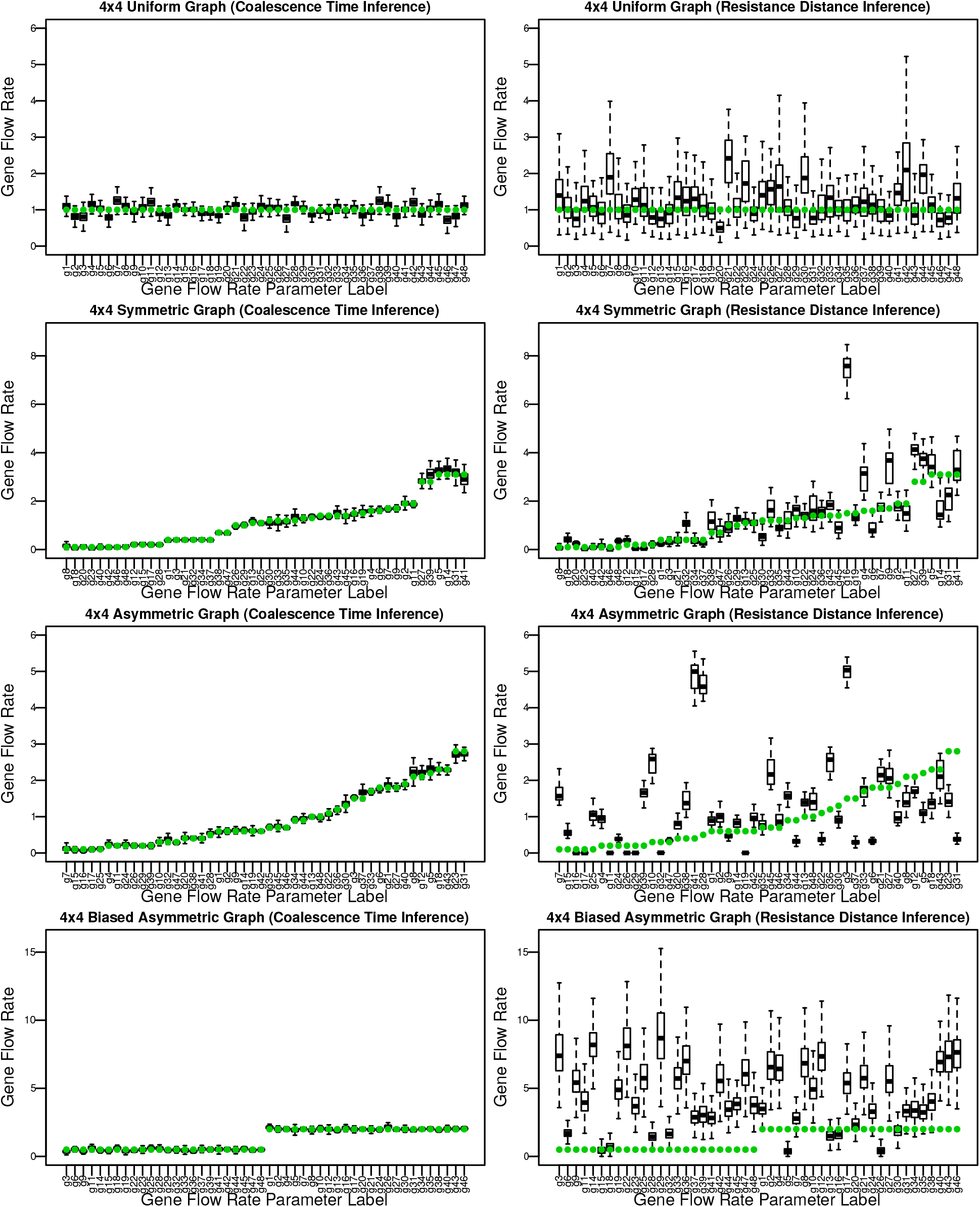
Boxplots of the posterior distributions of movement rates inferred using the **(left)** coalescence times and **(right)** resistance distances shown in Figure 4; i.e., on the 4×4 graphs with **(top)** uniform, **(middle top)** symmetric, **(middle bottom)** asymmetric, and **(bottom)** biased movement rates. See Figure S7 for specific values and graph structure.

**Figure 6:**
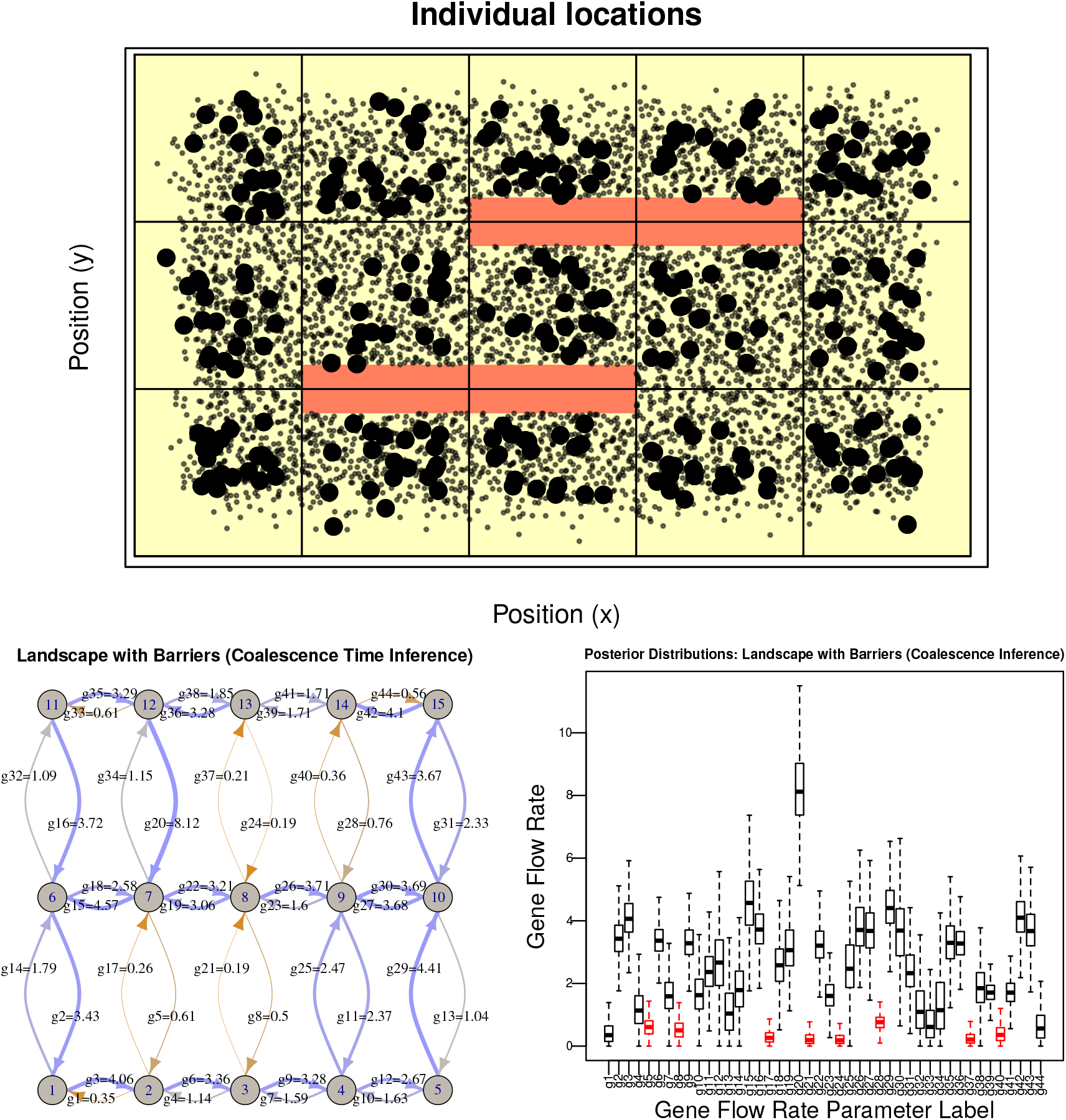
**(top)** Locations of individuals on the 5×3 landscape with barriers, with individuals used to compute the mean genetic distance matrix marked in bold. Red bars show uninhabitable regions, which are sufficiently thick so that no migration can occur directly across them. **(bottom left)** Posterior medians of movement rates inferred using coalescence time inference, and **(bottom right)** boxplots of corresponding posterior distributions, with movement rates crossing the barriers colored in red. The grouping of individuals into a 5 *×* 3 grid is shown above, and labels are the same as in Figure 2.

Each diploid individual has one pair of homologous chromosomes with 10^8^ loci each and a recombination rate of 10^*-*8^ per generation per locus. Neutral mutations were added at a rate of 10^*-*9^ per meiosis. To make these simulations computationally feasible, the mutation rate was set to 0.0 during the forwards simulation, while tree sequences were recorded [**?**], and mutations were added after the fact with msprime, which is equivalent to including mutations during the forwards simulation but resulted in much faster run times [Kelleher et al., 2018].

To compute genetic distances, geographic space was partitioned into square regions, and within each region, individuals are sampled uniformly for “genotyping” from among those within the middle three quarters of each dimension of the square, as shown for one situation in Figure 6. This protocol was chosen as a compromise between sampling all individuals from near the center of each location, which could result in a large number of close relatives being sampled, and uniform sampling from the whole area of each grid, which may result in many sampled individuals being very close to other grid squares. Mean genetic distances for each pair of regions are then computed as described above. To estimate uncertainty, we computed the standard error of these mean distances using the minimum number of individuals in each location.

Before being passed to the MCMC inference method, we rescaled genetic distances so that the movement rates resulting from the inference would be approximately of order one. We did this by multiplying genetic distances by a constant so that the rescaled mean of the entries of *D* is equal to the number of locations in the discretization.

### Data from *Populus trichocarpa*

We also applied our method to genotyping data from 423 samples of *Populus trichocarpa* and 8 samples of the closely related *Populus balsamifera* (with which it hybridizes), from a large region of northwestern North America, described in Moreno Geraldes et al. [2014b]. These data include geographical coordinates and genotypes from a genotyping chip targeting 34K SNPs. In addition to preliminary filtering described in Moreno Geraldes et al. [2014b], we additionally removed 2,314 SNPs with more than 5% missing data, retaining 30,756 SNPs, and two individuals, one with more than 15% missing data and one that is much further east than all the other samples. We then computed pairwise divergence between the remaining samples. We discretized the landscape manually into nine regions, preserving species boundaries and choosing divisions along gaps in sampling to keep geographic areas roughly equal. The numbers of samples included in each discretized region varied from 1 to 268, but our method does not assume equal sample numbers (however, an *F*_*ST*_ -based method would likely be affected by such large differences in sampling).

### Code and data availability

All code used for this work is available at https://github.com/elundgre/gene-flow-inference (including an R package that implements the MCMC method) and https://github.com/petrelharp/isolation_by_coalescence (including the SLiM simulations). Sequencing data for *Populus* is available from Data Dryad [Moreno Geraldes et al., 2014a].

## Results

### Model validation

We first tested the method using data generated under the model. To test the methods across a wide range of situations, we generated fifty sets of random movement rates on 3 *×* 2 rectangular grids: twenty-five symmetric (i.e., with *G*_*ij*_ = *G*_*ji*_) and twenty-five asymmetric. We generated movement parameters by rounding independent exponential random variables with mean 1 up to the nearest one tenth, and fixed coalescence rates at 1 for all locations. We then analytically computed mean coalescence times, and added independent noise to each pairwise coalescence time. We assume that we can estimate the amount of noise reasonably well, so used the true standard deviation of the noise for *σ*_*ϵ*_ in the likelihood function. In practice, *σ*_*ϵ*_ can be estimated by the standard deviation of genetic distances between different pairs of samples in the same region. To assess the impact on inference of the number of parameters, we performed inference both with and without the assumption that coalescence rate is the same everywhere. For 3×2 graphs, there are 21 equations and 14 movement parameters, so that with a global coalescence rate there are 15 unknowns. However, with individual coalescence rates there are 20 unknowns, which is close to nonidentifiable. To quantify accuracy across these random graphs in different situations, we computed, for each model fit on a particular random graph, the absolute difference between the posterior median of each movement parameter and the true value. We then averaged these across movement parameters to produce a measure of accuracy for that model fit, reported below as “mean absolute error”.

#### Varying noise

Unsurprisingly, coalescence time inference became less accurate as the amount of noise increased. As we varied the standard deviation of the noise from 1/1000^th^ to 1/50^th^ of the mean value of *C* for that grid, the mean absolute error of inferred gene flow rates (*G*_*ij*_) for coalescence time inference with a single coalescence rate increased from around 0.05 to 0.4. Since true values of *G*_*ij*_ were of order 1, this is a transition from precise to very rough (but still informative) inference. Posterior interquartile ranges increased proportionally, although there was substantial variation in accuracy between replicates; see Figures S2 and S3. Increasing the number of parameters by allowing multiple coalescence rates made the inference problem much harder – mean absolute errors no longer depended strongly on the amount of noise added, and hovered around 0.4.

More surprisingly, the performance of resistance distance inference did not depend on the amount of noise – mean absolute error was around 0.6 across all levels of noise, both with and without more than one parameter for local diversity (see Figure S2). Although symmetry did not affect coalescence time inference, resistance distance inference did substantially worse on asymmetric graphs, showing median absolute errors of around 0.8 – inferred gene flow rates were almost uncorrelated with true values.

#### Varying coalescence rates

Lower coalescence rates mean that lineages wander about the landscape for longer before they coalesce, thus greatly reducing the geographic information we get from coalescence times [Wilkins, 2004]. For instance, if between-population gene flow and coalescence rates are similar, then the lineages of two samples in the same population are likely to coalesce before either leaves, leading to a strong signal of isolation by distance. On the other hand, if coalescence rates are much smaller than gene flow, nearby samples are unlikely to share more recent common ancestors than distant ones. Indeed, coalescence rates had a strong effect on feasibility of the inference problem (shown in Figures S4 and S5): lowering the coalescence rate from 10 to 0.1 (always adding noise with standard deviation 1/200^th^ of the mean value) increased mean absolute error of coalescence time inference from around 0.05 to 0.5. (Gene flow rates are of order 1, so this range of coalescence rates spans strong to weak isolation by distance.) Again, allowing multiple coalescence rates or using resistance distance inference were highly inaccurate in all cases, with mean absolute errors of above 0.5.

### Identifying a barrier to gene flow

We then test the method’s ability to locate barriers to migration on the 5×3 grid with asymmetric gene flow shown in Figure 2. Each nonzero movement rate is determined randomly as before, and coalescence rates are set to 1 everywhere. Figure 3 shows posterior distributions of the movement rates for both coalescence time and resistance distance inference on this graph, with noise standard deviation equal to 1/1000^th^ of the mean coalescence time (top) and 1/100^th^ of the mean coalescence time (bottom). Although increased noise in the data increased uncertainty in the estimated values, coalescence time inference correctly inferred not only the location of the barrier but also the migration rates along all other edges within a reasonable margin of error. Resistance distance inference also identified the barrier, but was much less accurate for the other values. We also explored making the problem harder, by first (a) dropping data corresponding to populations 2, 5, 7, and 15; and then (b) allowing a separate coalescence rate for each location, in both cases using a noise level of 1/100. The resulting posterior distributions are shown in Figure S6, and show substantially increased error, but only in movement rates to the removed locations. As before, uncertainty was increased when multiple coalescence rates were allowed, but not as strongly as before, likely because the problem is less close to being underconstrained, having 105 equations and 59 unknowns.

### Biased migration

Both matrices of coalescence and commute times, *C* and *R*, are symmetric, but they do not deal with asymmetric (biased) migration in the same way. Since commute time to location *i* requires paths both to and from *i*, but coalescence time to *i* does not, if there is a low rate of migration back into *i*, then commute times to *i* could be quite long even while coalescence times are not. For instance, suppose there are three populations arranged in a line, and lineages currently in the outer populations are much more likely to come from the central population than the other way around. Since lineages quickly move to the center and coalesce regardless of starting position, coalescence time (and genetic diversity) will be relatively low between all individuals. Commute time between the two outer locations, on the other hand, will be much longer than other comparisons, since it requires a lineage to leave the center.

In order to investigate this general situation, we tested both methods on four different 4×4 graphs (shown in Figure S7): *uniform*, where all movement rates are 1.0; *symmetric*, where movement rates are symmetric (*G*_*ij*_ = *G*_*ji*_) and randomly generated as before; *asymmetric*, where all movement rates are random; and *biased*, where movement rates down or to the left are equal to 2.0, and movement rates up or to the right are 0.5. For inference, we added noise with standard deviation 1*/*500^th^ of the mean value.

Differences between true mean coalescence times (*C*) and mean resistance distances 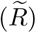 are shown in Figure 4: they are fairly small in the uniform and symmetric cases, moderate in the asymmetric case, and extreme in the biased asymmetric case. This difference suggests that inferences made with resistance distance may be strongly misleading in the asymmetric and biased situations, but in principle it still could be the case that the movement rates that give the best fit of resistance distance to coalescence time data could be close to the actual rates use to generate the data. This does not seem to occur in practice: coalescence time inference was substantially more accurate in all cases. Posterior distributions of inferred movement parameters are shown in Figure 5. The mean absolute errors for coalescence time inference are 0.12 or less in all cases. Resistance distance inference obtains roughly uniform migration rates (although varying by a factor of 2) in the uniform case; and migration rates noisily correlated with the truth in the symmetric graph. However, resistance distance inferences for the asymmetric and biased graphs are only weakly correlated, if at all, to the truth. Resistance distance inference also drastically overestimates movement rates for the biased case, as we would expect based on the differences discussed above.

### Continuous geographical space

The data we have used thus far are produced under the model used for inference, and should provide an accurate depiction of our method applied to discrete, randomly mating populations whose connections by migration are known. However, this is nearly always a rough approximation to reality, in which organisms are distributed across continuous geography, and “populations” are constructed by necessity, often driven by sampling locations. For this reason, we applied coalescence and resistance inference, as well as the resistance-based method EEMS [Petkova et al., 2016], to data from simulations on continuous geography.

#### Process noise

The mean coalescence times of lineages that we compute analytically are exact for large, randomly mating populations, but the word “mean” indicates an average across several levels of randomness: each observed, empirical mean estimated using a sample of genotypes will deviate from this depending on the individuals sampled (“individual sampling noise”) [Ashander et al., 2018], the genetic loci genotyped (“genome sampling noise”) [Waples and Faulkner, 2009], and the stochasticity of the population history itself (“process noise”) [Wakeley et al., 2012, Waples and Faulkner, 2009]. Sequencing of many nonascertained loci genome-wide should minimize the second source of noise (although genome structure can affect this at large scales [Li and Ralph, 2019]). Geographic models in continuous space, are expected to show much more process noise than discrete population models because of random fluctuations in local population density. To quantify the relative contributions of the remaining sources of noise, we compared mean genetic distances, calculated in the same way, between two nonoverlapping sets of samples from each of three identical simulations. These forwards-time, continuous-space simulations were done using SLiM as described in the Methods. We found that with these parameter values, process and sampling noise were of similar magnitude, contributing 6.7% and 13.8% of the variance, respectively (Supplemental Figure S8).

#### Biased migration

To test inference on a simple, continuous landscape with bias, we simulated a square, flat landscape as above, but with *biased migration*: each offspring’s location was chosen randomly as before, but with mean location *σ*_*d*_*/*10 up and to the right of the parent’s location. This again produces reverse-time gene flow down and to the left. Coalescence time inference correctly infers the bias on a broad scale: using a 4×4 grid, inferred gene flow rates down and to the left are consistently much larger than those up and to the right; resistance distance-based results show no clear pattern (Supplemental Figure S10). However, we do not obtain the clear picture of the landscape we saw for the corresponding discrete-population model (bottom-right plot, Figure 5): inferred rates show substantial heterogeneity, likely due to stochasticity of recent demography.

#### Identifying a barrier

We now revisit the earlier situation where a landscape has locations that are barriers to gene flow. The landscape is shown in Figure 6, where red bars depict uninhabitable regions that block migration. Since the distribution of offspring locations is truncated at three standard deviations away from the mother, a sufficiently thick uninhabitable area completely stops migration directly across it. We calculate mean genetic distances using 50 individuals from each grid location, here giving us a total of 750 individuals. The gene flow rates across the barriers are reliably estimated to be very small, while other values have substantial variation: in particular, gene flow rates are estimated to be low across some edges that do not cross barriers; this may be because of recent demographic stochasticity. The barriers were also identified with comparable accuracy by both our resistance distance method and EEMS, as shown in Supplemental Figure S11. Note also that this analysis began by grouping samples into discrete “populations” appropriately; in absence of a good *a priori* idea of where barriers are, this step might not be straightforward. To make parameters most easily interpretable, geography should probably be discretized along natural barriers and so that each region is of roughly the same geographic area.

#### Other barriers: population sinks and recent expansions

We simulated two other situations on a continuous landscape that produced an bias in effective migration. First, we simulated a **valley** in a square landscape, in which fecundity of individuals in the middle 1/4 of the landscape was reduced by 50%. The valley is a population “sink”, although it would not be detectable with mortality rates, and shows only a slightly lower population density (Supplemental Figure S12). Since individuals in the valley are more likely to have parents outside of the valley, this produces gene flow “uphill”, away from the ridge. Next, we simulated a recent **expansion**, in which the middle 4/5 of the landscape was uninhabitable for 400,000 generations, until 500 generations before the present day. (This is a model of secondary contact with a long period of isolation, but not outside the realm of possibility.) The resulting expansion produces gene flow back away from the center, similar to the “valley”.

Both coalescence time inference and the resistance landscape inferred by EEMS show a barrier to gene flow in the middle of the landscape (Figure 7). This is arguably correct, although there is in fact no current barrier to migration in either simulation. Coalescence time inference also correctly infers the general bias in recent gene flow away from the center, although as in other continuous-space simulations, there was substantial noise in the inferred patterns. However, the resistance landscapes as inferred by EEMS show patchy, vertical bands of high resistance. These patterns are commonly seen when attempting to use EEMS, whose underlying model of gene flow is symmetric, to describe anisotropic gene flow. The grid-like pattern visible in this and other EEMS results may be due to the sampling scheme (recall that samples are drawn from the middle three-quarters of each grid square). Our analysis above shows that this is an unavoidable consequence of the resistance approximation, and our implementation of resistance distance inference, which allows asymmetric gene flow, shows similar patterns (Supplemental Figures S12 and S13).

**Figure 7:**
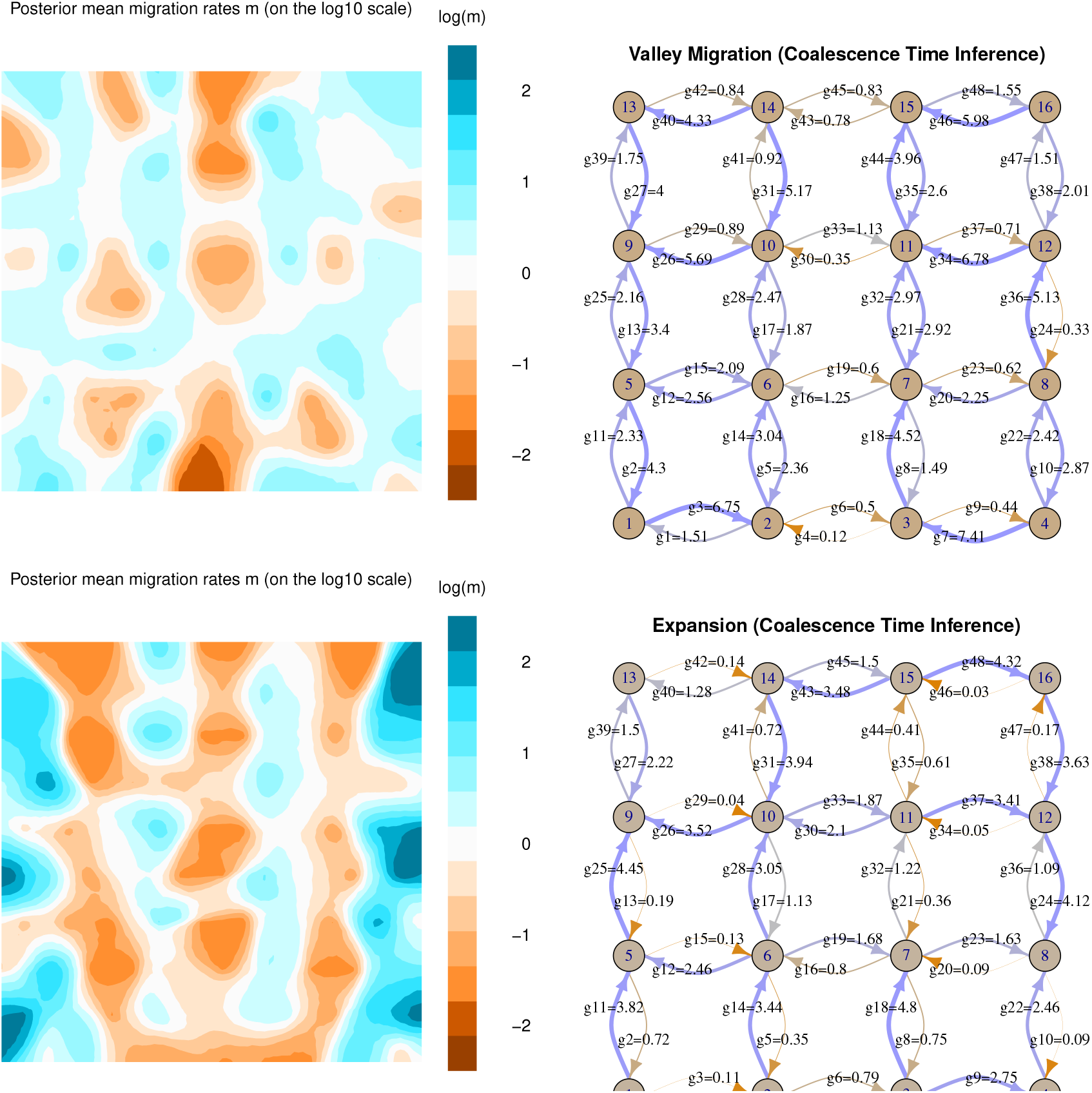
**(Left:)** resistance landscapes inferred by EEMS and **(right:)** gene flow rates inferred using coalescence time inference, from individual-based simulations on a continuous landscape, with **(top)** a *valley* of reduced fecundity in the middle, and **(bottom)** a recent *population expansion* from both edges of the range that met in the middle. The color scale shows the posterior mean log migration rates inferred by EEMS.

### Application to *Populus*

Our discretization of the sampling area for the two poplar species is shown in Figure 8, along with arrows depicting posterior mean estimates of gene flow. Posterior distributions of gene flow rates and coalescence rates are shown in Supplemental Figures S14 and S15. The main features of the resulting model are: (a) low but nonzero gene flow between species, with the highest rate between regions 3 and 4; (b) strong bias in gene flow among (inland) *balsamifera* regions, from southeast to northwest (3→5→6→1); and (c) strong bias in gene flow into regions 2 and 8. These results align with what is known about *Populus* history and ecology in the region (personal communication, A. Moreno Geraldes and Q. Cronk, and Moreno Geraldes et al. [2014b]). There is ongoing hybridization around region 4, and interspecific gene flow from 9 to 3 may be due to downstream transport along the Columbia river (a larger gene flow rate from 9 to 3 indicates that individuals in region 9 are more likely to have recent ancestry in region 3 than vice-versa.) The glacial refugia for the two species are thought to be in western Alaska (*balsamifera*) and coastal British Columbia (*trichocarpa*). Inferred gene flow leading back towards these regions (depicted by larger arrows on the map) are consistent with this – for instance, if region 5 was colonized from region 6 sufficiently recently, then individuals in region 5 should trace their ancestry to region 6 much more frequently than vice-versa, as indicated by the large 5→6 arrow and small 6→5 arrow in Figure 8.

**Figure 8:**
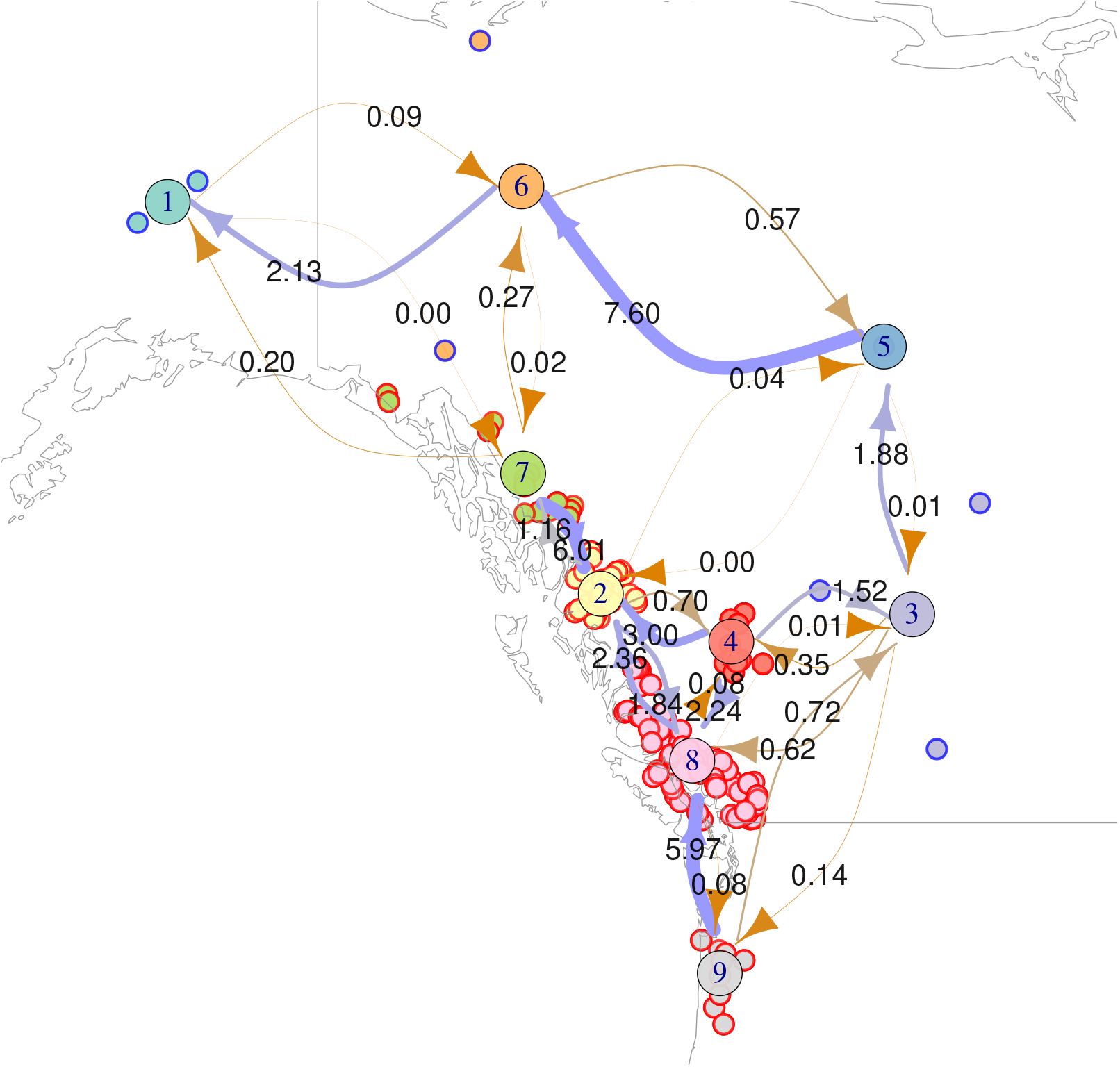
Model results for *Populus* data, superimposed on a map of northwestern North America. Sampling locations of *Populus* genomes are small circles, with outer color indicating species (red-outlined, coastal samples are *trichocarpa*, blue-outlined, inland samples are *balsamifera*); circle fill color indicates how samples are grouped into discrete “populations”. Large circles are placed at group centroids, and given group labels. Arrows are labeled by the posterior mean gene flow rate inferred under the coalescence time model (which also determines arrow width and color). For instance, the inferred gene flow rate from 4 to 2 is 3.0, while that from 2 to 4 is 0.7, suggesting that poplars in region 4 have many more recent ancestors in region 2 than poplars in region 2 do in region 4. Northwestern bias in gene flow between *balsamifera* regions may reflect postglacial expansion to the south and east out of a refugium in Alaska.

Interpreting these results requires some nuance beyond what was necessary in our simulation results. Sampling locations are sparse, especially in the inland species, so how exactly should we think of the geographic extent of “region 6”, which is represented by only two, distant samples? Is this region’s low inferred coalescence rate a result of simply a large population (due to a large geographic region) or barriers to movement within the region? Although it is not entirely clear how to translate discrete-population models to continuous space, a “gene flow rate” from region *i* to region *j* as shown in Figure 8 is best thought of as an estimate of the proportion of the individuals in region *i* that have a parent in region *j*, averaged over time and scaled by some unknown factor. Equivalently, this is the probability per unit time that a lineage traced back from an individual in region *i* follows a line of descent to region *j*, again scaled by some factor. We therefore expect gene flow to be biased from an area of lower population density (or, net fecundity) into a neighboring region of higher population density. We also expect gene flow to be biased from a smaller region to a larger, neighboring one than vice-versa if a larger proportion of the smaller region lies close to the boundary between the two. This may explain the net bias from many coastal populations to inland ones. We grouped samples so that each group represented roughly comparable geographic regions, to ameliorate these confounding factors. However, these difficulties in interpretation are common in any situation where discrete populations are used to model continuous geography.

## Discussion

In this paper, we study how to use genetic and geographic distances between present-day samples to infer the demographic parameters of a population that lives across a heterogeneous two-dimensional landscape. In particular, we have shown that the resistance distance approximation – which underlies several of the most commonly-used tools of landscape genetics – can produce erroneous estimates, especially in the presence of biased gene flow. We implement an alternative method, which uses coalescence times instead of the resistance distance approximation, which provides good estimates of gene flow in a wider range of situations (Figures 3 and 5). This weakness of resistance-based methods is not surprising – the original papers describing Circuitscape [McRae, 2006] and EEMS [Petkova et al., 2016] present the resistance approximation as a necessary step for computational feasibility. Results of resistance-based methods should therefore be interpreted with caution: for example, high inferred gene flow between two areas may actually mean the areas are being fed by a common source.

Our new method infers effective population sizes and gene flow rates quite well – and, being Bayesian, provides estimates of uncertainty – given data from discrete populations connected by migration. Can our method replace resistance-based methods? Perhaps, but the substantial uncertainty we saw on graphs with only tens of nodes is indicative of a larger problem we face for realistic models. We have seen that discretization of space results in substantial modeling error, in part because of randomness of geographic sampling and unmodeled process noise can lead to overfitting. Partitioning space into a finer grid should help with these problems, but tends to make the inference problem itself more ill-conditioned: with more connections, changing the value of one connection affects coalescence time less, and so inferences about that value must necessarily be less certain. This tradeoff implies some degree of unavoidable uncertainty. Reproducible, reliable inference will likely require development of new inference methods that explicitly model continuous geography. In the meantime, comparison of results utilizing different discretizations of geography can help identify problems.

Although commute and coalescence times can be quite different, they are equal (with a particular choice of local diversity) if populations are arranged on a ring or torus with symmetric migration (as also noted by McRae [2006]). The similarity of these models to rectangular grids may explain why in other work, resistance distance methods have shown relatively good fit to data simulated with symmetric migration rates.

### Other methods

For comparison on equal footing, we have implemented a resistance-based inference method. However, methods that use resistance distance have substantial advantages over the method we present here. Circuitscape [McRae et al., 2008] produces maps of much higher resolution than is presently possible with coalescent times. Hanks and Hooten [2013] connects the problem to intrinsic conditional autoregressive models, providing a Bayesian model of an underlying Gaussian Markov random field. EEMS [Petkova et al., 2016] integrates over possible tessellations of the region as a way of regularizing the inferred landscape, aiming to obtain a map at the best possible resolution supported by the data.

Most promisingly, the recent method, MAPS [“Migration and Population Surface estimation”, Al-Asadi et al., 2018] is based on EEMS, but instead of resistance distances, uses pairwise sharing of long haplotype segments, which have been shown to carry substantial information about recent demography [Ralph and Coop, 2013, Palamara and Pe’er, 2013, Browning and Browning, 2015, Ringbauer et al., 2017]. Furthermore, the likelihood model underlying MAPS is based on coalescent theory, and thus should not be subject to the drawbacks of resistance distance. Ringbauer et al. [2018] also uses shared haplotype lengths to identify a barrier in a coalescence-based framework. However, the underlying migration model is symmetric, and haplotype length-based methods require fairly dense genotyping and a genetic map, something not available for many species.

Other widely-used inference methods, such as MIGRATE [Beerli and Felsenstein, 1999, Beerli and Palczewski, 2010], BayesAss [Wilson and Rannala, 2003], and IMa [Hey and Nielsen, 2007] use likelihoods for the joint frequency spectrum derived from a coalescent model – and so in principle could use much more of the data than our method, which only uses pairwise divergences. However, because they use locus-specific data rather than an average across loci, these methods can only be applied to a relatively small number of populations and loci, because of computational efficiency. We have found asymmetry in migration rates to be important: Hanks [2017] developed a method that allows asymmetry by modeling genetic similarity as deriving from an underlying random field whose space-time covariance is given by the covariance of the (forwards-time) population fluctuations. However, this is not motivated by a generative model for genetic data.

All of the methods presented above work with a discretized model of space, for mathematical and computational reasons. Development of new methods for continuous geography and assessment of how current methods behave when faced with continuous geography is an important challenge facing the field. This has been made substantially easier with the introduction of continuous space into the simulator, SLiM [Haller and Messer, 2018], that we used to test the behavior of our method with more realistic data. Another promising approach centers around the Spatial Lambda-Fleming-Viot model [Barton et al., 2010], which provides a mechanistic model in which coalescent simulations are possible. This has been used for inference [Guindon et al., 2016], but more work remains to understand how it behaves as an approximation to real populations.

### Asymmetric dispersal and consequences for resistance methods

We have found that asymmetric rates of lineage movement can produce qualitative differences between coalescence times and resistance distances, and thereby mislead models based on resistance distance. How should this affect the results of resistance-based methods to genomic data, besides general inaccuracy? This asymmetry occurs if one region receives a greater proportion of each new generation as migrants from another population than that other population does from it. This is expected, for instance, if organismal dispersal is biased by wind or water currents [Gaines et al., 2003, Morrissey and de Kerckhove, 2009], or in the presence of source-sink dynamics [Dias, 1996, Lenormand, 2002]. Suppose, for instance, that seed dispersal for a species of tree is biased downhill, and the landscape is dominated by a large valley. Since lineages tend to move uphill, genetic distance between locations on opposite slopes of the valley will be relatively large, so resistance methods will infer a barrier along the bottom of the valley when no barrier to dispersal exists. On the other hand, if there is a ridge instead of a valley, resistance methods will show no barrier (and relatively high movement rates), even if there is very little uphill dispersal. This could lead to a falsely high assessment of gene flow across the top of the ridge.

### Circuitscape

Could we use coalescence time in place of commute time for workflows that compute values on a given map of movement rates that are then compared to genetic distance? In principle this could be done, but there is no clear way to do this at the same (impressively fine) geographic resolution that is currently possible, e.g., using Circuitscape [McRae, 2006]. This is because if we have *n* sampled locations on a landscape discretized into *N* regions, the computational complexity of finding commute times is *nN*, but coalescence time scales as *N* ^2^ (Each *N* -vector of hitting times to a particular location can be found independently, but the entire *N*×*N* matrix of coalescence times must be found together.) However, as we discuss above, this fine geographic resolution is in some sense illusory – for each fine-resolution map there should be coarser maps that give almost identical coalescence times – but we are not aware of existing theory providing guidance on how to find these.

### Difficulty of inference

What determines the feasibility of inferring movement rates from genetic distances? Using coalescence times instead of commute times greatly improves accuracy in many situations, but there are still a number of factors that determine the tractability of the problem. Observation noise is clearly an issue: we found that estimation errors of a few percent was enough to seriously degrade accuracy. However, genetic distances can be estimated to much higher accuracy using modern genomic data. Perhaps more seriously, a low rate of coalescence relative to movement can also make the problem essentially non-identifiable. This is clear at least in the limit: with very low coalescence, the population loses any isolation by distance and is indistinguishable from a randomly mating population. In continuous landscapes, this balance is measured by Wright’s local effective population size, which is proportional to the number of other individuals within a circle of radius equal to the mean dispersal distance.

Inference of movement rates, in either a coalescence time or resistance distance framework, is an ill-conditioned problem, implying the need for regularization to obtain reliable inference. This fact could explain the results of Graves et al. [2013], who found a large area of resistance parameters that produced equivalently good fits to genetic distance (i.e., nonidentifiability, analogous to a flat likelihood surface). An anonymous reviewer suggested that although inference of landscapes of movement rates may carry a degree of unavoidable uncertainty, the problem of *model comparison* (e.g., between models with and without migration across a certain barrier) may still produce more certain answers.

Computation time currently limits this method to a discretization of less than 30 populations, but nonidentifiability is probably a more serious barrier that must be concurrently addressed before scaling the method to finer discretizations of space. Finer discretizations of space should benefit from the use of sparse matrix methods, but in our testing these did not speed up computation at this scale.

### The effect of history

The models we discuss here assume that population sizes and migration rates have been constant on the time scale given by the within-species coalescent time. This is rarely true in practice [Neigel et al., 1991, Barton and Wilson, 1995]. However, geographic differences in mean relatedness are established on a shorter time scale – the time scale over which a lineage, moving randomly across the landscape, “forgets” where it started (i.e., the mixing time of the Markov chain [Wilkins, 2004]). This is the time it takes for the uncertainty in lineage movement to reach the scale of the species range. For instance, if the width of a species’ range is roughly 500km, and if mean dispersal distance is 10km, the standard deviation of ancestor location *t* generations ago is 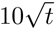, so the mixing time of a lineages across the landscape, not accounting for barriers, is of order 2500 generations. The habitat of many species would have changed substantially over this time, which suggests that models incorporating change over time may be required to model modern diversity. However, the effects of recent history are strongest, so landscapes estimated assuming constant populations may give a reasonable picture of the landscape averaged over recent times.

### Assumptions

Modeling lineages as a Markov chain is nearly ubiquitous in population genetics today, but may not be appropriate. Even if the population dynamics in forwards time are Markov, the dynamics of lineages traced back in time (the coalescent process) may not be. For example, if individual fecundity is variable and population density is low, two lineages near each other are more likely to share a recent common origin, so knowing how one lineage moves is informative about how the other lineage is likely to move. This effect becomes small as population density increases, so it should be a relatively minor point if the offspring of any one parent are typically interspersed with the offspring of many others. Natural selection also makes the coalescent process non-Markovian. Differences in the strength of linked selection along the genome could even cause the statistical behavior of lineages to depend on the region of the genome being studied [Wang and Bradburd, 2014, Li and Ralph, 2019]. However, more investigation with continuous-space models is needed.

## Acknowledgements

Thanks to David Levin for useful suggestions regarding hitting time calculations, to Paul Marjoram for useful comments, to Brad Shaffer and Evan McCartney-Melstad for discussions about landscape modeling, to John Novembre, Hussein Al-Asadi and Benjamin Peter for help with EEMS and general input, and to Quentin Cronk and Armando Moreno Geraldes for consultations about the *Populus* results. Thanks also go to five anonymous reviewers for numerous useful suggestions. We would also like to gratefully acknowledge the immense positive impact that Brad McRae [1966–2017; Lawler et al., 2018] has had on landscape genetics and conservation biology through the introduction of landscape resistance. We hope this paper helps to carry forward this work. Work on this project was supported by funding from the Sloan Foundation and the NSF (under DBI-1262645) to PR.

## A The simplest example

To demonstrate the main theoretical ideas, we provide a short example. Consider a Markov chain with two states, where the rate of movement from state 1 to state 2 is *G*_12_ and the rate of movement from state 2 to state 1 is *G*_21_, so the generator matrix is

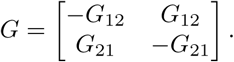

Equation 7 equates two 2*×*2 matrices, so provides four equations. Only three of these are unique; simplifying these and using that *C*_12_ = *C*_21_, these are:

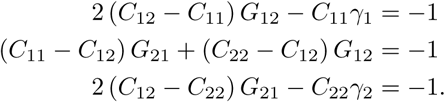

Given *C*, we have three equations for the four unknowns, *G*_12_, *G*_21_, *γ*_1_, and *γ*_2_, which we can solve symbolically. The valid solutions to these equations are those with *G*_12_, *G*_21_, *γ*_1_, and *γ*_2_ nonnegative. If *C*_11_ ≠ *C*_12_, we can write the solution with *γ*_1_ as the free variable:

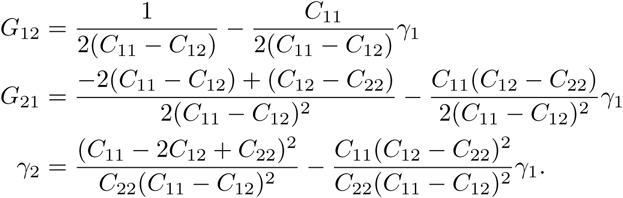

This implies that *γ*_2_ decreases as *γ*_1_ increases as long as *C*_12_ ≠*C*_22_. This makes sense because increasing a rate of coalescence cannot make expected coalescence times longer, so in order to keep coalescence times the same, if one coalescence rate is increased, another must be decreased. This also implies that, in order for *γ*_2_ to be a value other than 0, we must have *C*_11_ *-* 2*C*_12_ + *C*_22_ ≠ 0. In general, if there is a value or range of values of *γ*_1_ for which the other movement and coalescence parameters are non-negative, then a solution exists.

To produce a concrete example, suppose that *C*_11_ = 1, *C*_12_ = 2, and *C*_22_ = 1.5 in arbitrary time units. The other parameters will be non-negative when 1 ≤ *γ*_1_ ≤ 5, as shown in in Figure S1. Assuming *γ*_1_ = *γ*_2_ produces a unique solution, with *γ*_1_ = *γ*_2_ = 1.29 and *G*_12_ = 0.14 and *G*_21_ = 0.93.

## B Finding *G* from *H*

Suppose that *G* is the generator matrix of an irreducible continuous-time Markov chain *X* on at least two states, so that *G*_*ij*_ ≥ 0 for *i* ≠*j* and *G***1** = 0. Under these assumptions, there is a unique stationary distribution, *π*, that satisfies *π*^*T*^ **1** = 1 and *π*^*T*^ *G* = 0. Let *τ*_*j*_ = inf {*t* ≥0 : *X*_*t*_ = *j*} and *H*_*ij*_ = 𝔼 [*τ*_*j*_|*X*_0_ = *i*].

Then we know that

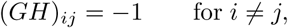

and hence that for some vector *x*,

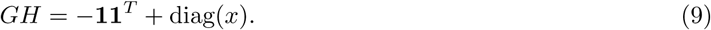

What is *x*? First note that the Random Target Lemma [Aldous and Fill, 2002] tells us that if *π* is the stationary distribution of the chain, then (*Hπ*)_*i*_ does not depend on *i*, and so there exists a vector *v* ∝*π* such that *Hv* = **1**. Multiplying equation (9) by *v*, we get that

**Figure S1:**
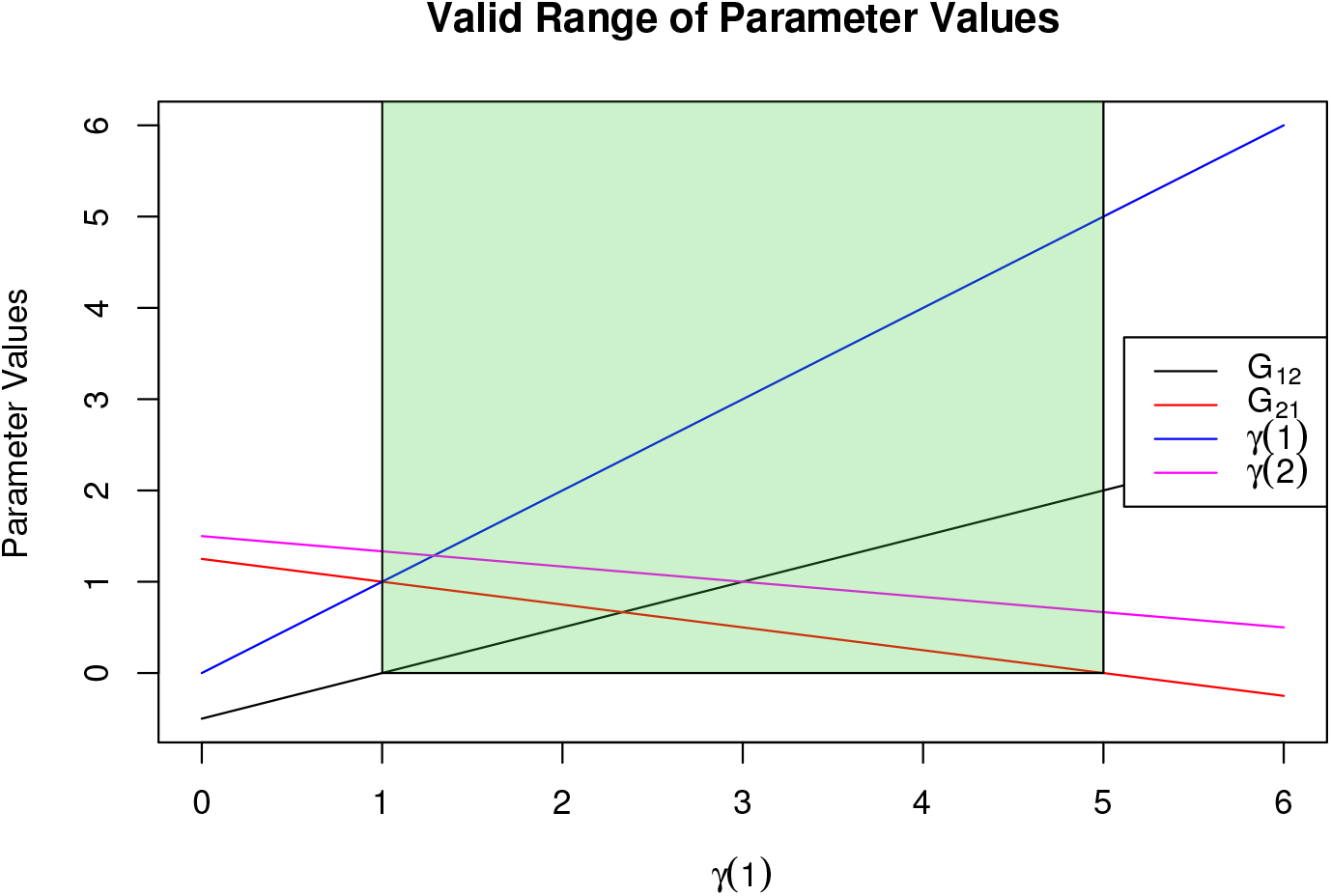
The green shaded region shows the valid range of the parameter values for the two state Markov chain given that *C*_11_ = 1, *C*_12_ = 2, and *C*_22_ = 1.5

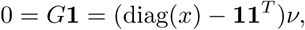

which rearranges to

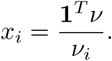

But, since ν ∝ *π*, it must be that *π* = ν */***1**^*T*^ ν, and so *x*_*i*_ = 1*/π*_*i*_.

Now, we prove that *H* is in fact invertible, by showing that *z*^*T*^ *H* ≠0 for every nonzero vector *z*. First note that **1**^*T*^ *H* ≠0, since the entries of *H* are nonnegative. Now take *z* orthogonal to **1**. Since the chain is irreducible, the eigenspace associated with the eigenvalue 0 for *G* has multiplicity one, and so any vector *y* that satisfies *y*^*T*^ *G* = 0 is proportional to *π*. This implies that there is a vector *y* such that *y*^*T*^ *G* = *z*^*T*^. Multiplying equation (9) on the left by *y*^*T*^ obtains *z*^*T*^ *H* = *y*^*T*^ (diag(1*/π*) *-* **11**^*T*^). If *H* is not invertible, then there is a *z* for which this is zero, i.e., *y*_*i*_*/π*_*i*_ = Σ_*j*_ *y*_*j*_ for every *i*. But, this only holds if *y* = *π*, which was disallowed, because then we would have *z* = 0. Therefore, *H* is invertible.

In summary, *π* = *H*^*-*1^**1***/***1**^*T*^ *H*^*-*1^**1**, and

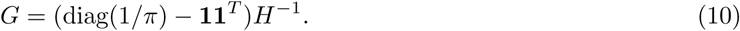

Note that this equation only specifies *H* up to a constant added to each column: i.e., if given *G* one obtains a matrix *Y* solving *GY* = diag(1*/π*) *-* **11**^*T*^, then *H*_*ij*_ = *Y*_*ij*_ *- Y*_*jj*_.

## C Equality of commute and coalescence times

Under what conditions are coalescence times and commute times (with some local diversity values) equal? In other words, for what choices of *G, γ*, and *q* does

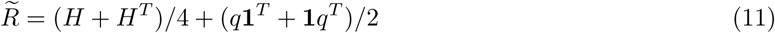

solve equation (7)? Writing this out, this says that

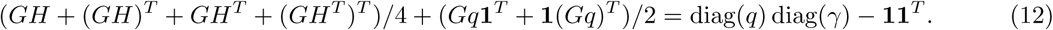

(To simplify this we used the fact that diag(*H*) = 0 and *G***1** = 0.)

It is not clear what can be said about the general case because of the presence of *HG*^*T*^, but if we assume that *H* is symmetric, we can make progress, because then we have that *GH* = *GH*^*T*^, and so equation (9) says that all four terms in the first group are equal, and equation (12) simplifies to

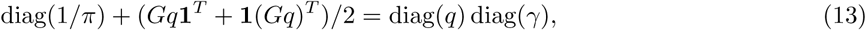

where the -**11**^*T*^ terms on each side canceled. Because *G***1** = 0, the most obvious solution to this is if *q* = *c***1** and *cγ* = 1*/π*, for some constant *c* (although there is a broader family of solutions).

In summary, if hitting times are symmetric and coalescence rates are equal to the inverse of the stationary distribution, then coalescence times are equal to commute times plus one. This is fairly restrictive, but does occur if the population configuration is isotropic (as for instance in an all-connected-to-all island model with equal migration rates) or in populations arranged around a ring with migration rates depending only on the distance between them.

Can we solve equation (13) more generally? For that equation to hold, we need *Gq***1**^*T*^ + **1**(*Gq*)^*T*^ to be diagonal, i.e., that (*Gq*)_*i*_ + (*Gq*)_*j*_ = 0 for all *i ≠ j*. For any *k≠* ℓ, there exists a vector *u*^(*kℓ*)^ such that (*Gu*^(*kℓ*)^)_*i*_ = *δ*_*ik*_ *-δ*_*if*_; the only possible *q* for which *Gq***1**^*T*^ + **1**(*Gq*)^*T*^ is diagonal are of the form *q* = *α***1**+ *βu*^(*k, ℓ*)^ for some *k≠ℓ* and some constants *α* and *β*. We would then need

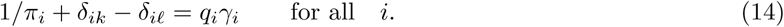

This implies that for any Markov chain with symmetric hitting times, we can find diversity values (*q*) and coalescence rates (*γ*) that make 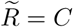(and in fact there are many ways to do this). However, there is not a general solution for *q* if coalescence rates are also given (as is the case in practice). This is related to the fact proved by Strobeck [1987] (and generalized by Nagylaki [1998]) that diag(*C*) is constant and does not depend on movement rates for any isotropic conservative migration model.

**Figure S2:**
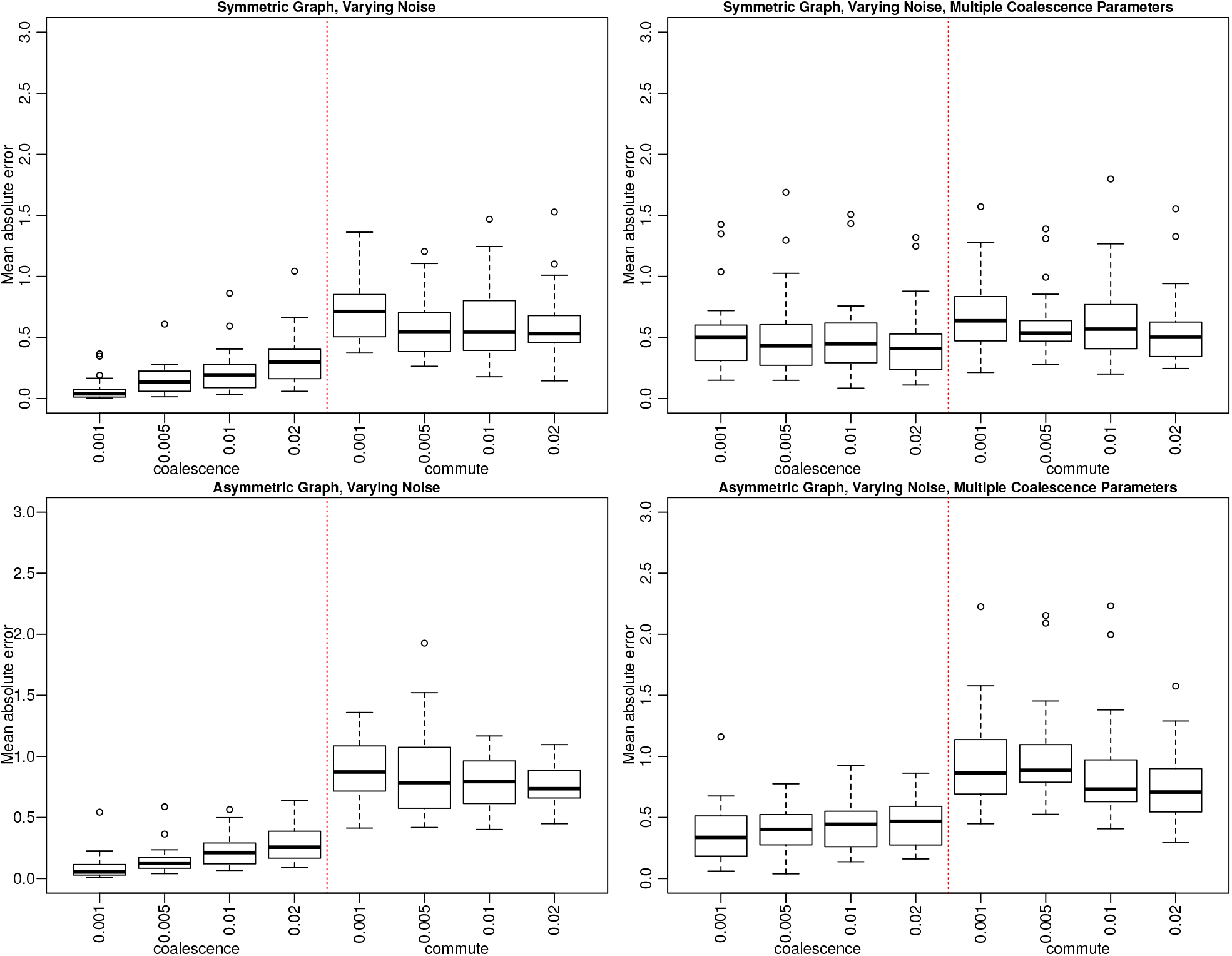
Boxplots of the mean absolute error defined to be the absolute value difference between the true value and posterior median, averaged across movement parameters *g* for each of the 25 graphs in each situation. Each box is labeled with the relative amount of noise added to *C* to produce the values used for inference. The top left shows the results for symmetric graphs when there is a single coalescence parameter for all locations, the top right for symmetric graphs when there is a separate coalescence parameter for each location, the bottom left for asymmetric graphs when there is a single coalescence parameter for all locations, and the bottom right for asymmetric graphs when there is a separate coalescence parameter for each location.

**Figure S3:**
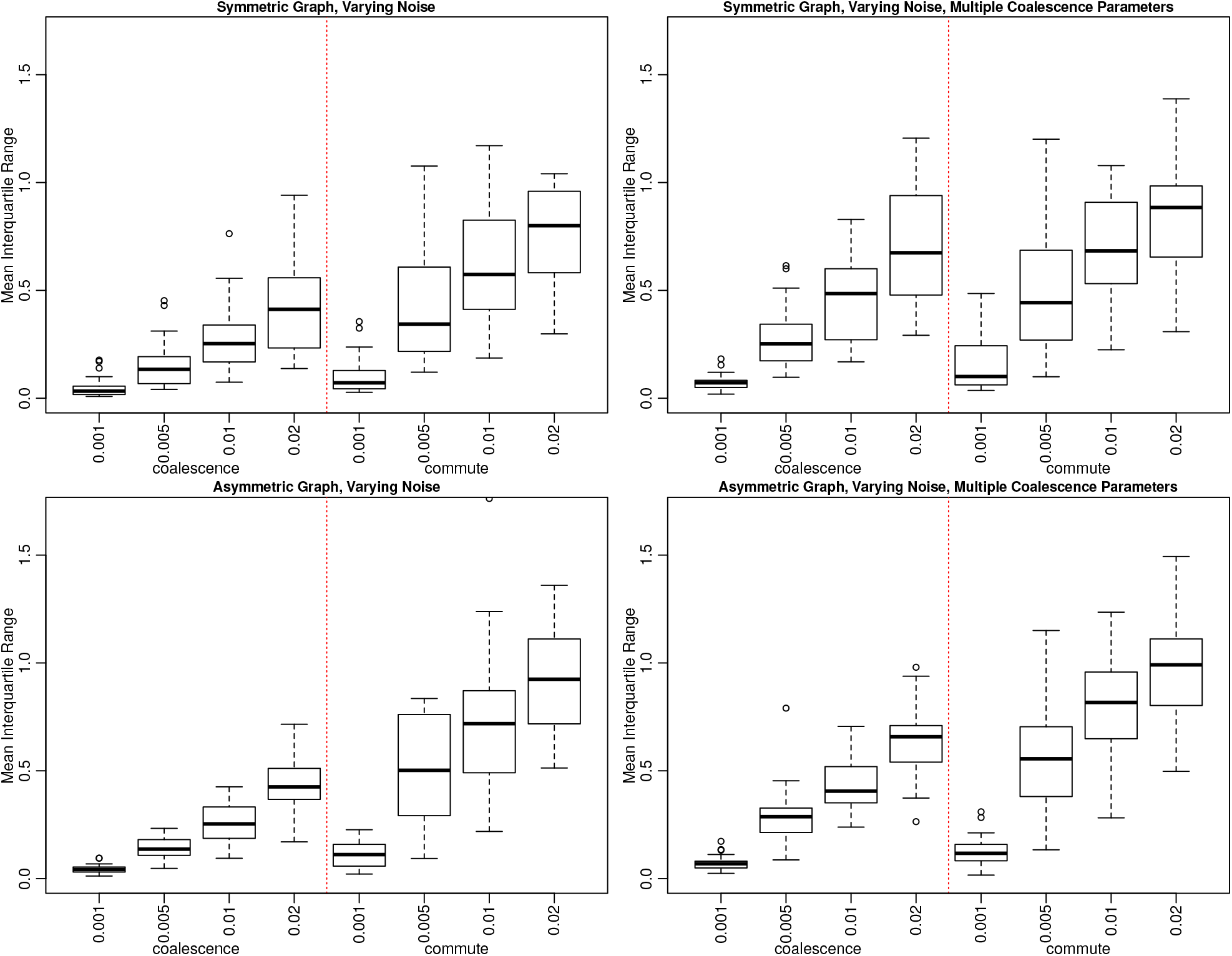
Boxplots of the mean interquartile ranges of the posterior distributions of *g* for each of the 25 graphs in each situation. Each box is labeled with the relative amount of noise added to *C* to produce the values used for inference. Subplot locations for each situation are the same as in Figure S2.

**Figure S4:**
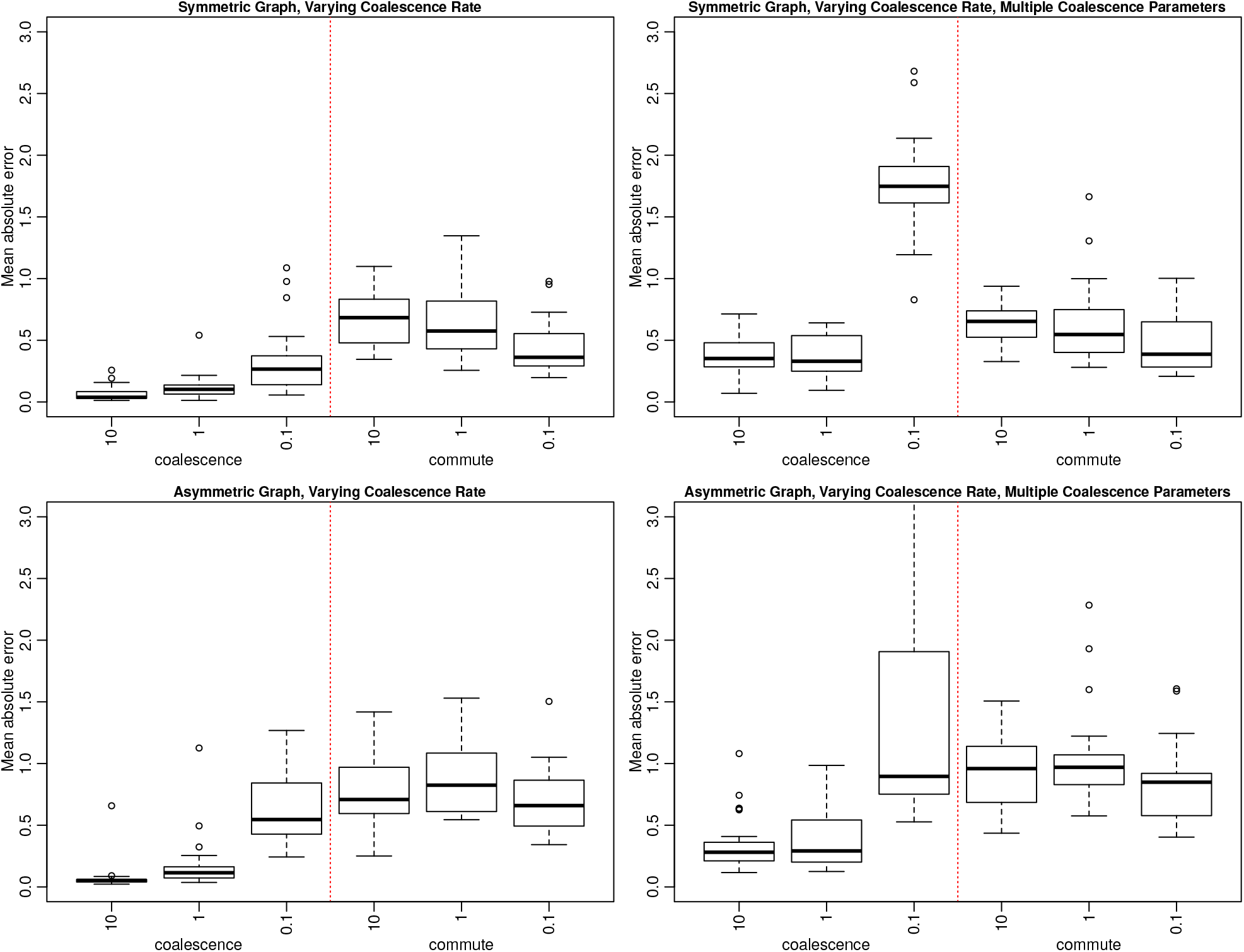
Boxplots of the mean absolute error, defined to be the absolute value difference between the true value and posterior median, averaged across movement parameters *g* for each of the 25 graphs in each situation. Each box is labeled with the coalescence rate in the underlying model for each situation. Subplot locations for each situation are the same as in Figure S2.

**Figure S5:**
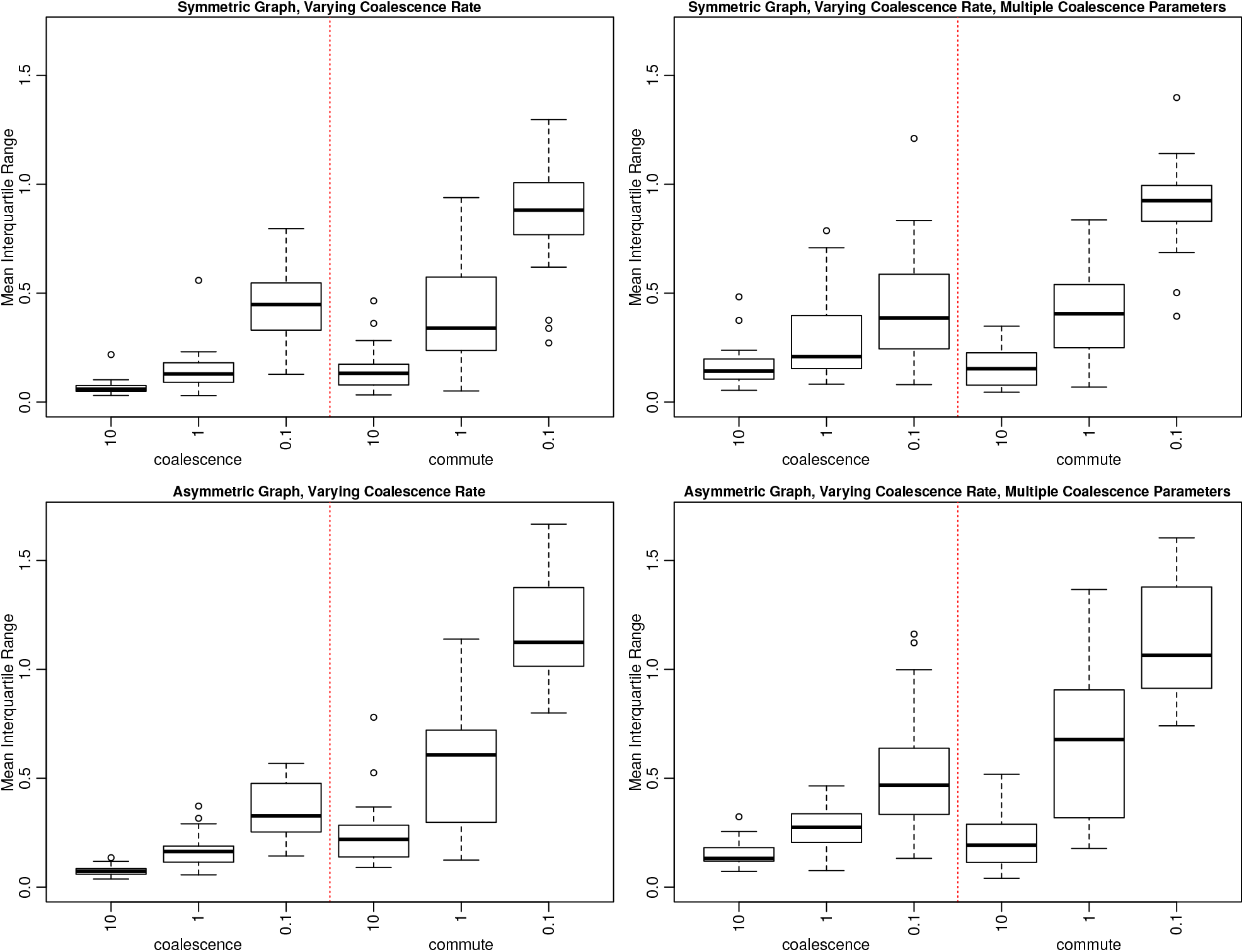
Boxplots of the mean interquartile ranges of the posterior distributions of *g* for each of the 25 graphs in each situation. Each box is labeled with the coalescence rate in the underlying model for each situation. Subplot locations for each situation are the same as in Figure S2.

**Figure S6:**
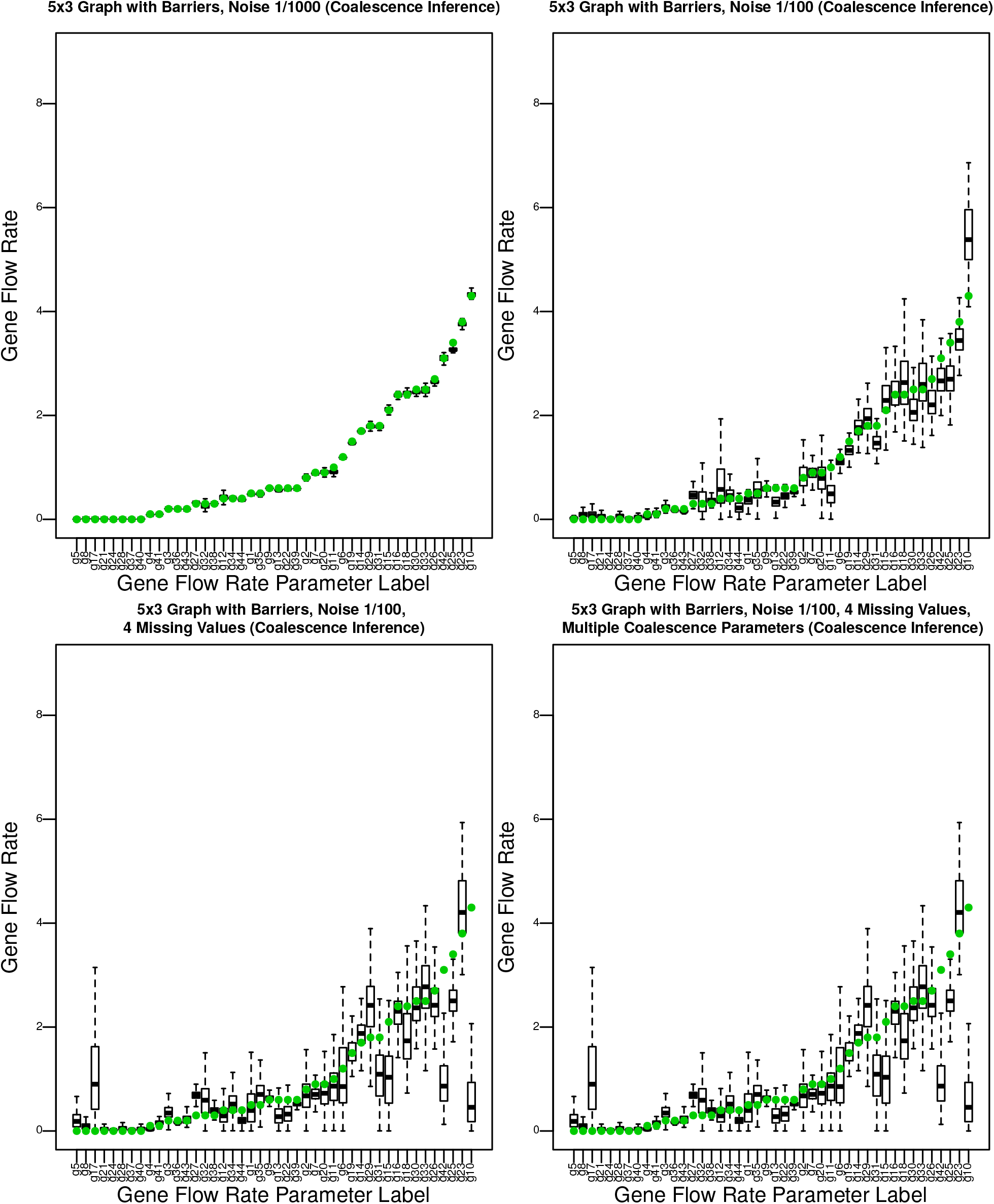
Posterior distributions for values of *g* for the 5×3 graph with barriers for each of the analysis cases with coalescence time inference. See Figure 2 to see movement parameter locations.

**Figure S7:**
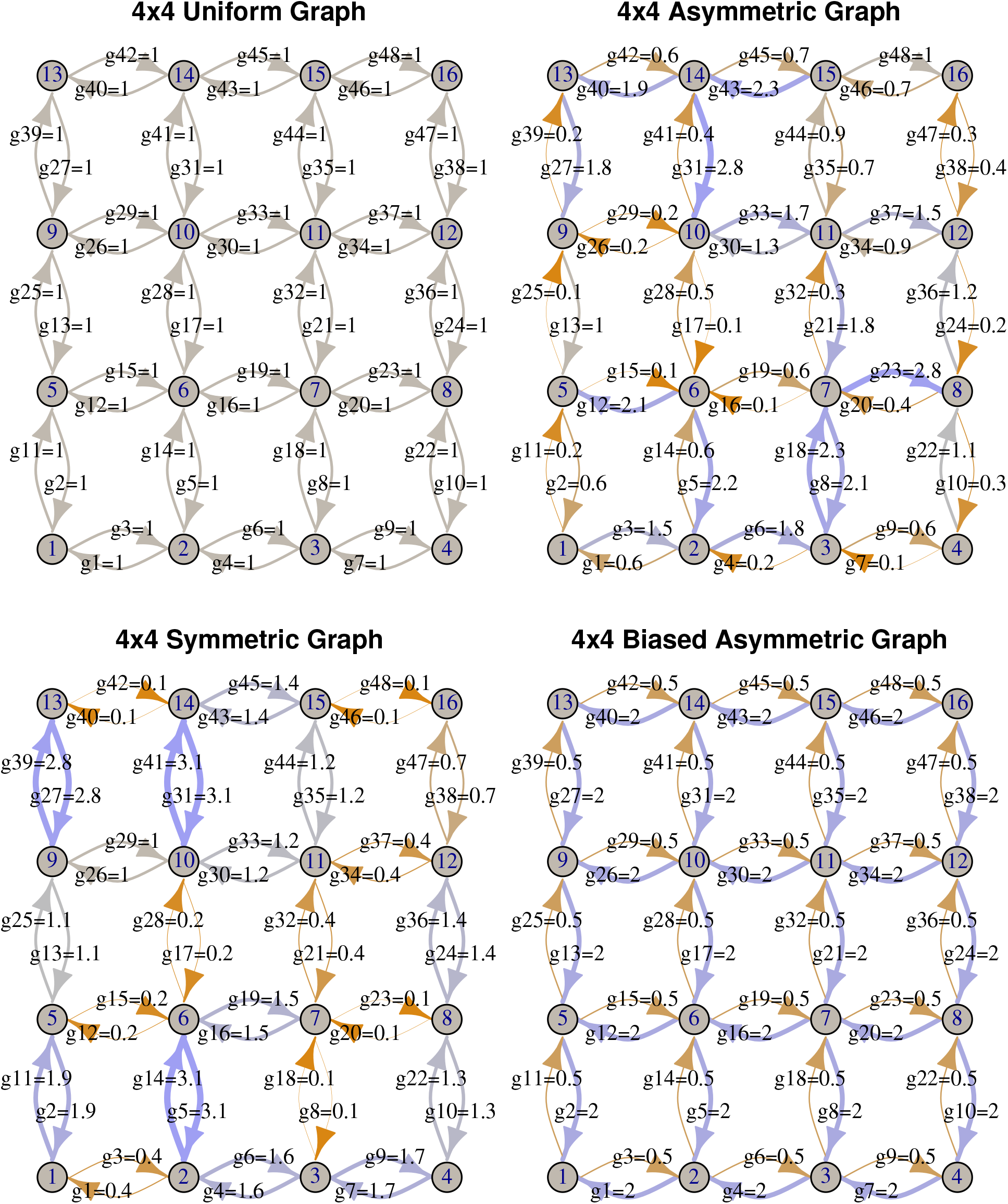
Grid structure and gene flow rates for the uniform graph, symmetric graph, the asymmetric graph, and the biased asymmetric graph using the igraph R package.

**Figure S8:**
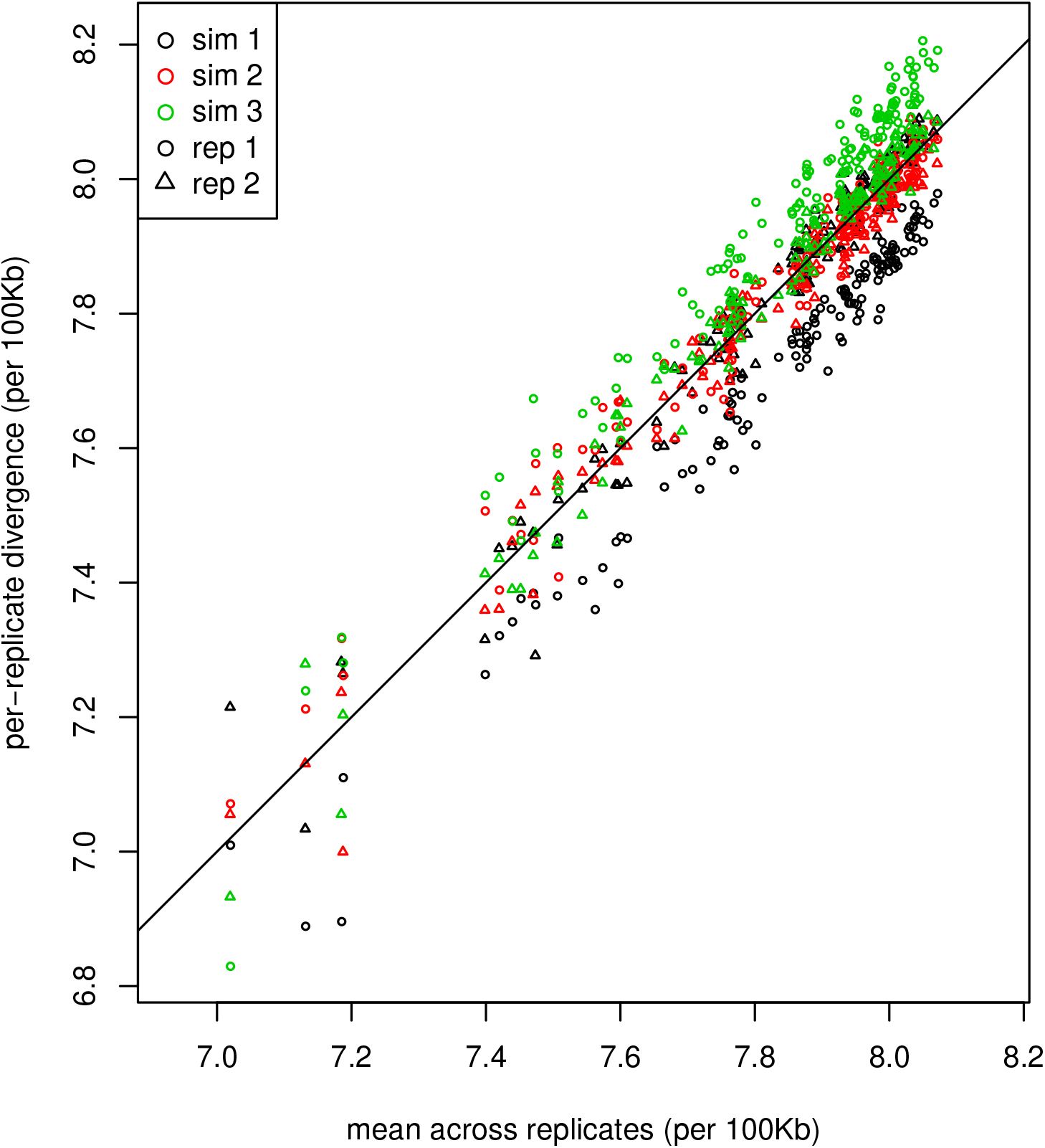
Comparison of mean coalescence times between samples and between replicates for a 4×4 discretization of a square landscape with uniform migration. Between the 16 geographic regions, there are 136 comparisons; these are shown for two nonoverlapping samples taken from three independent replicate simulations, each plotted against the mean value across all replicates. Colors denote simulation replicate, and point type (circle/triangle) denotes the sample. Without sampling or process noise, all points would fall on the *y* = *x* line. Among the six sets of observations, simulation replicate explains 6.7% of the variance, and spatial sampling explains 13.8% of the variance.

**Figure S9:**
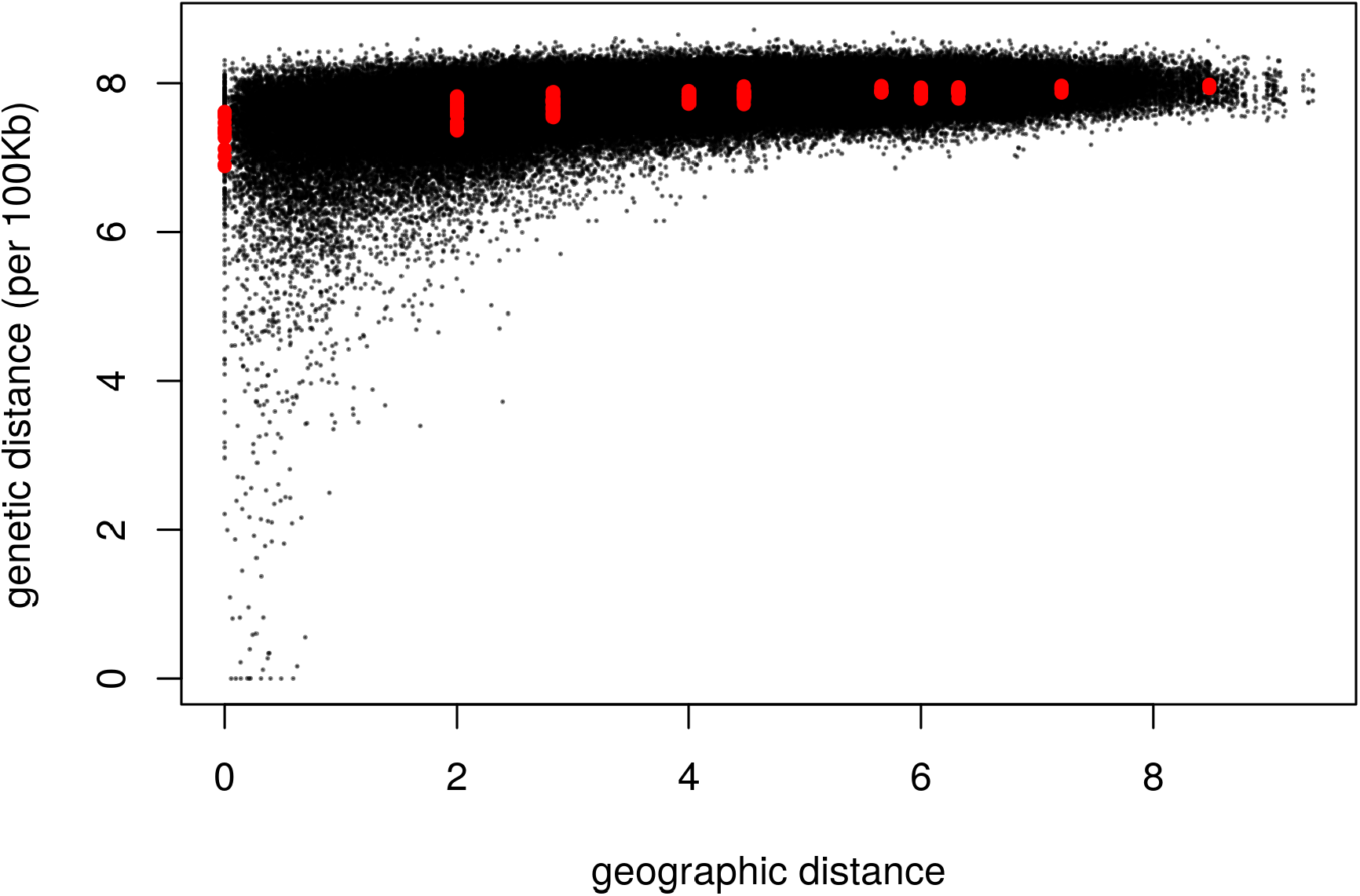
Genetic distance against geographic distance between all pairs of sampled individuals on a square landscape with uniform migration. Red dots denote mean genetic distances between sampled individuals from a 4 *×* 4 discretization of the landscape, as used for instance in Figure 7.

**Figure S10:**
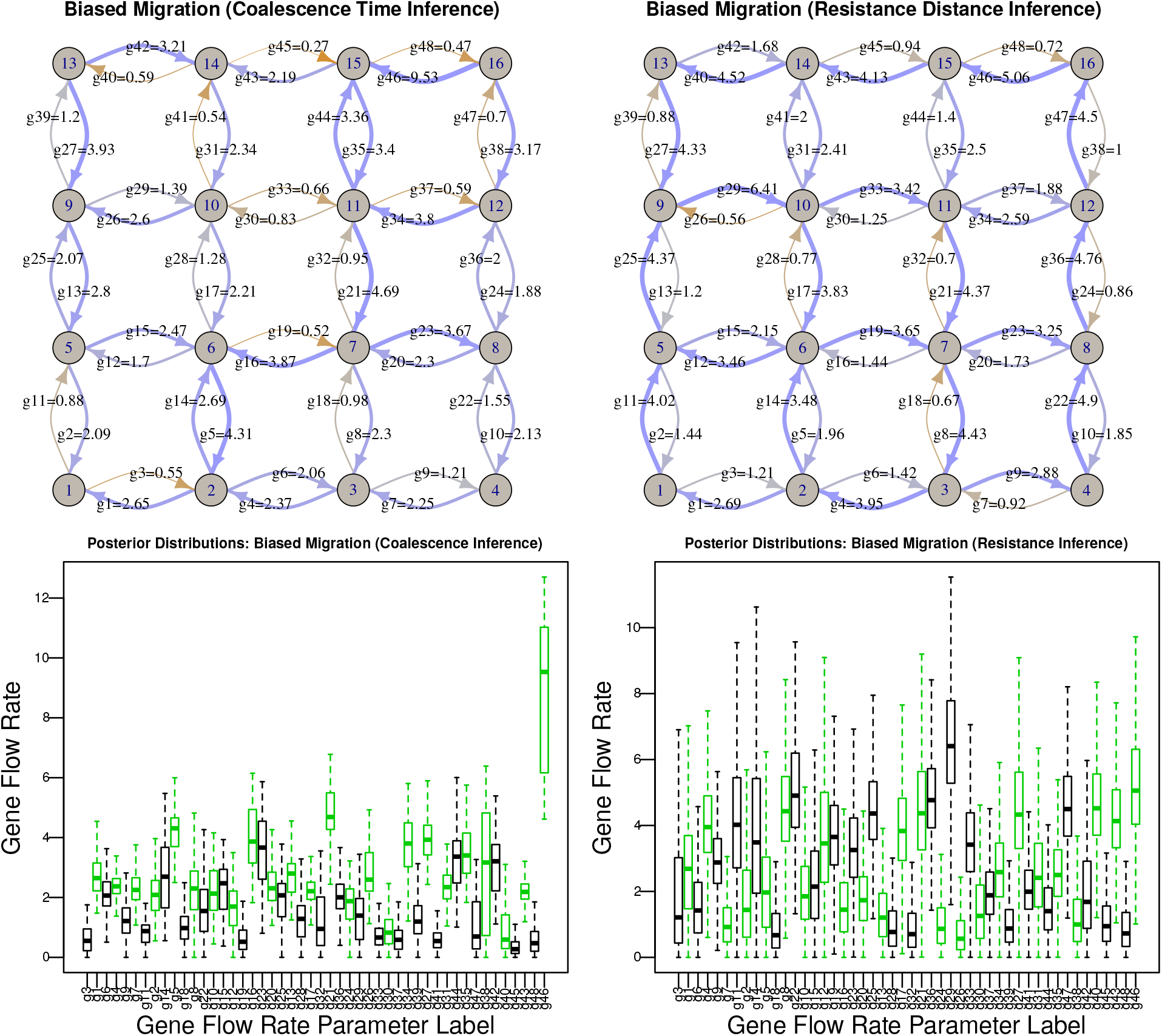
**(Above:)** Posterior medians and **(below)** posterior distributions of the inferred values of *g* for the 4×4 discretization of the square landscape with biased migration, inferred with the **(left)** coalescence time-based, and **(right)** resistance distance-based method. In the boxplots, black boxes show posterior distributions of gene flow rates up and to the right, and the adjacent green box corresponds to gene flow rate along the same edge in the opposite direction.

**Figure S11:**
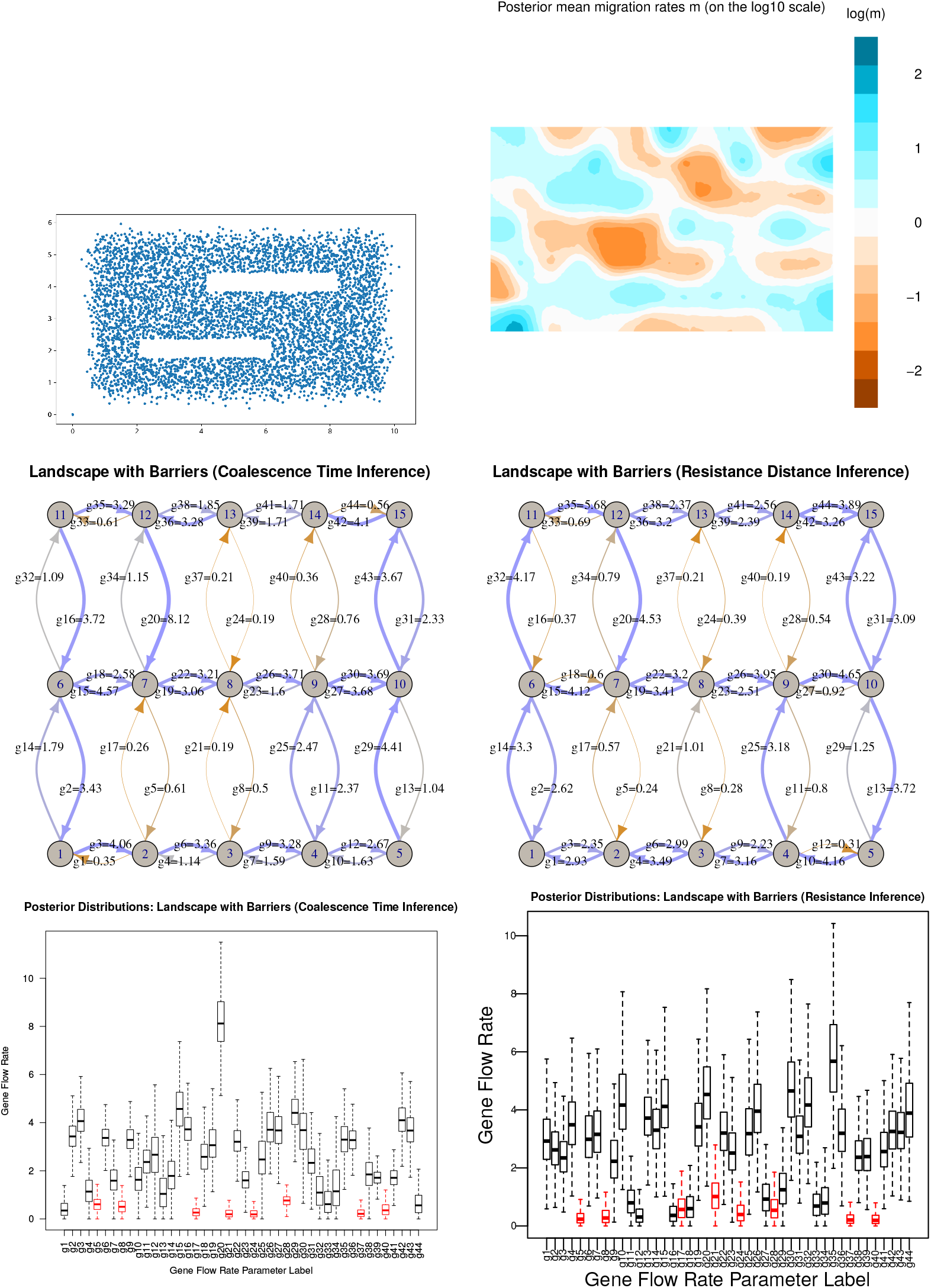
Results from the **barrier** simulation: **(top left)** Positions of all individuals at the end of simulation. **(top right)** mean migration rates as estimated by EEMS. Below are shown posterior median migration rates as estimated using **(middle left)** the coalescence time method and **(middle right)** the resistance distance method. Last are shown the posterior distributions, for **(bottom left)** the coalescence time method and **(bottom right)** the resistance distance method.

**Figure S12:**
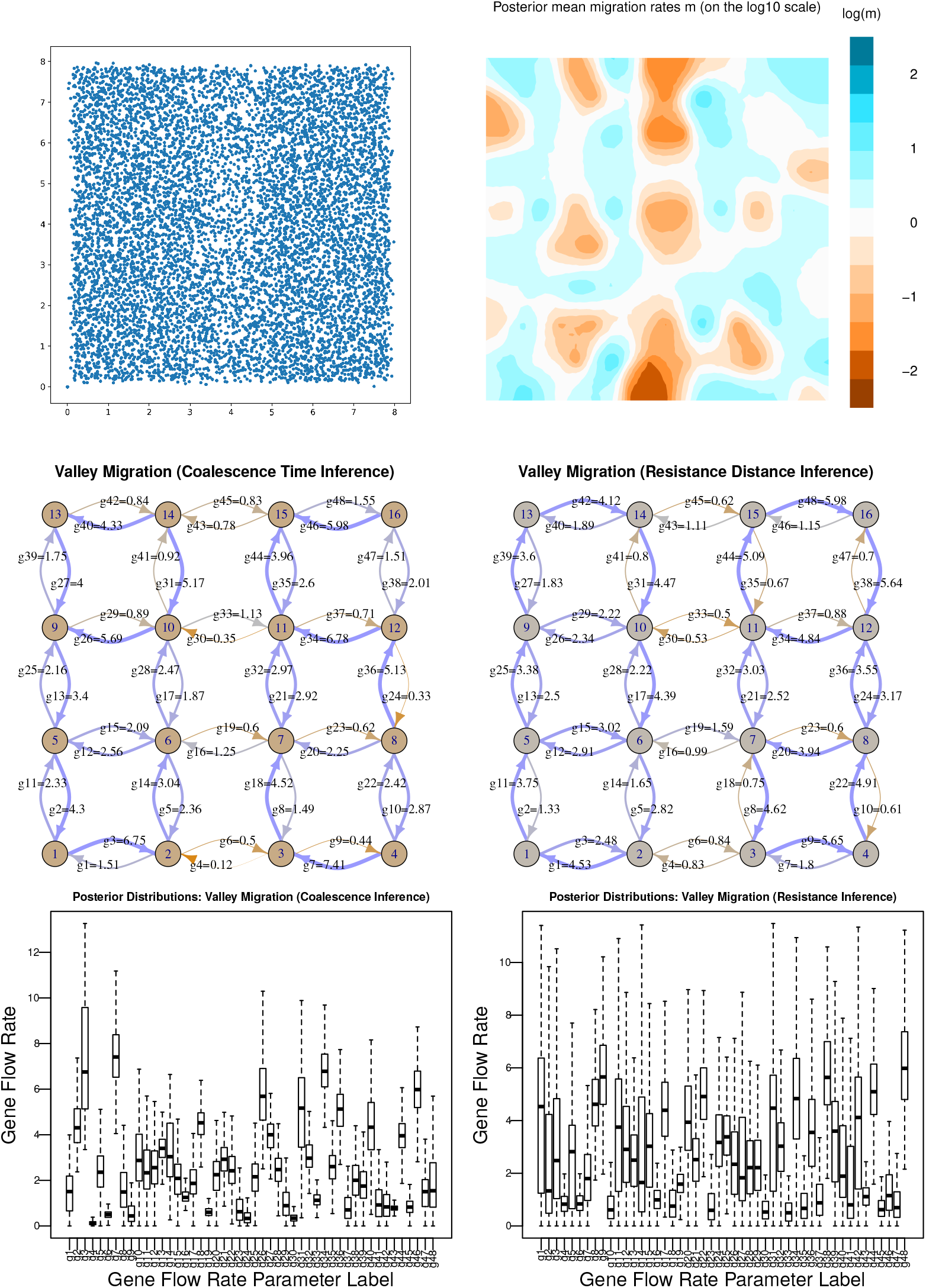
Results from the **valley** simulation: **(top left)** Positions of all individuals at the end of simulation. **(top right)** mean migration rates as estimated by EEMS. Below are shown posterior median migration rates as estimated using **(middle left)** the coalescence time method and **(middle right)** the resistance distance method. Last are shown the posterior distributions, for **(bottom left)** the coalescence time method and **(bottom right)** the resistance distance method.

**Figure S13:**
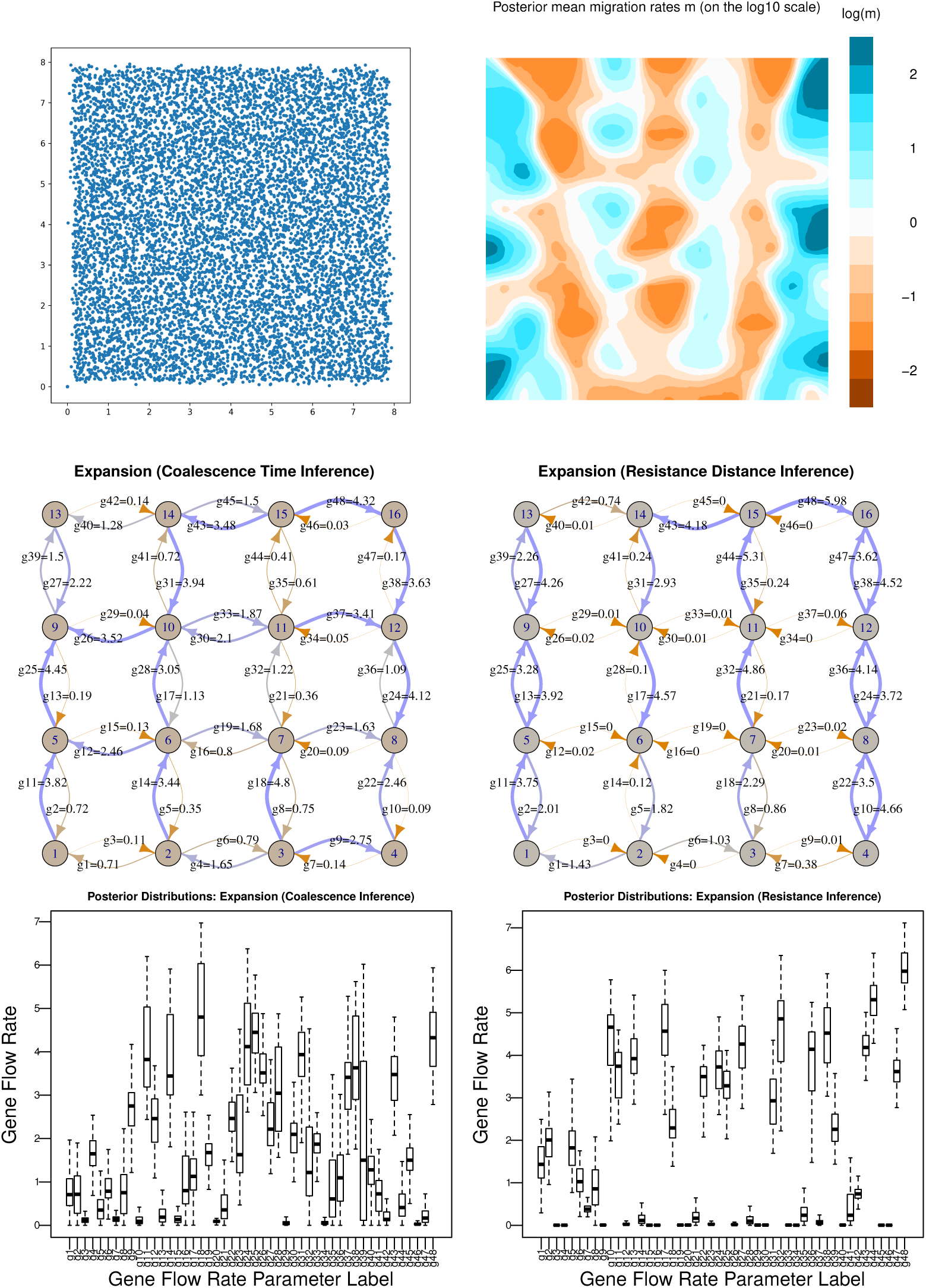
Results from the **expansion** simulation: **(top left)** Positions of all individuals at the end of simulation. **(top right)** mean migration rates as estimated by EEMS. Below are shown posterior median migration rates as estimated using **(middle left)** the coalescence time method and **(middle right)** the resistance distance method. Last are shown the posterior distributions, for **(bottom left)** the coalescence time method and **(bottom right)** the resistance distance method.

**Figure S14:**
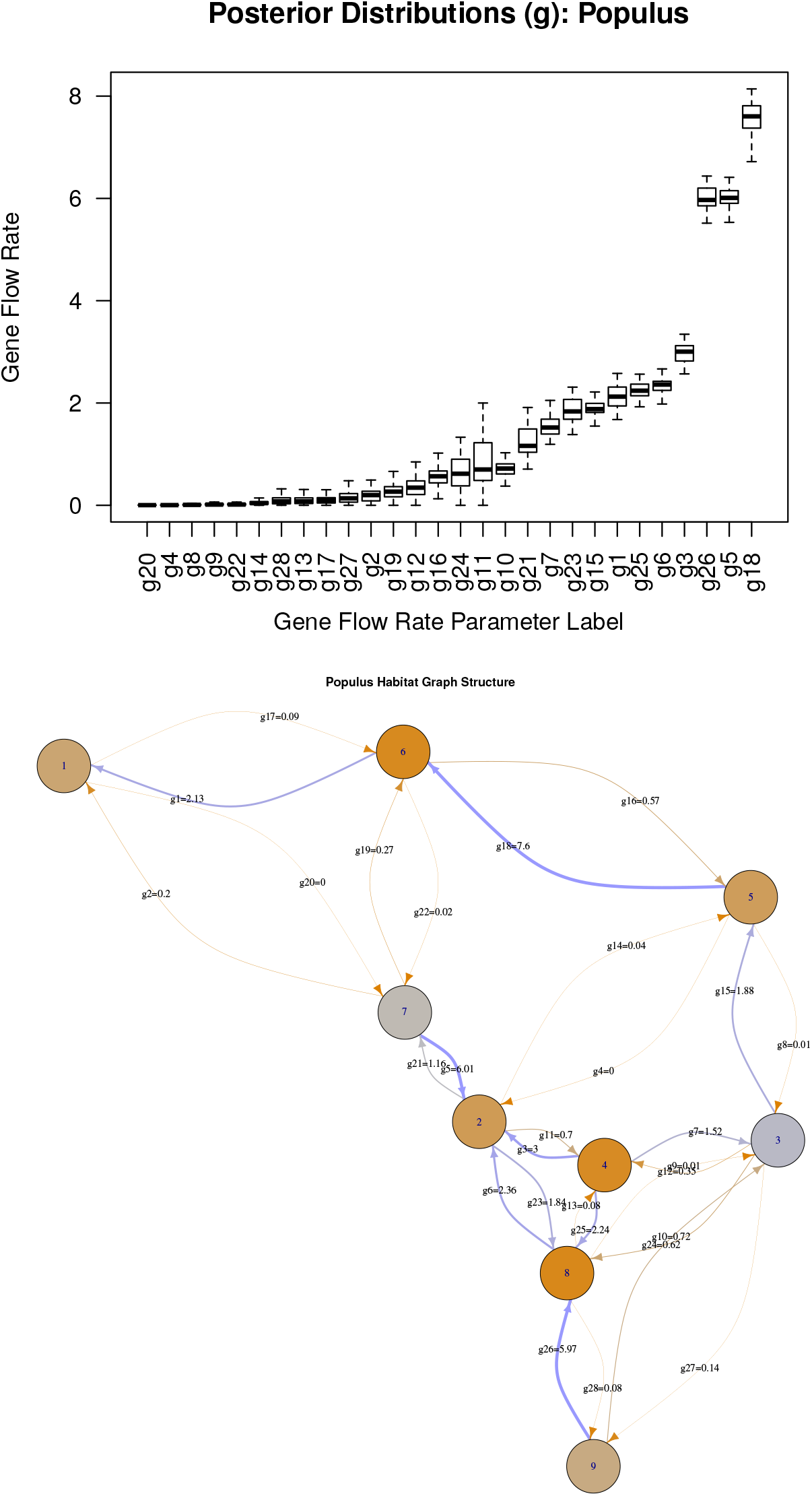
Posterior distributions of parameters for *Populus* data, using coalescence time inference with the model shown in Figure 8. **(top)** Posterior medians (dark line), approximate middle 50% (box), and 90% (whiskers), for gene flow rates. **(bottom)** Posterior medians gene flow rates, both as a numerical label and color (as in other figures, red is lower and blue is larger), along with edge labels that match the upper figure. Nodes are also colored according to posterior median coalescence rate: grey is a high rate (small population) and orange is a low rate (large population).

**Figure S15:**
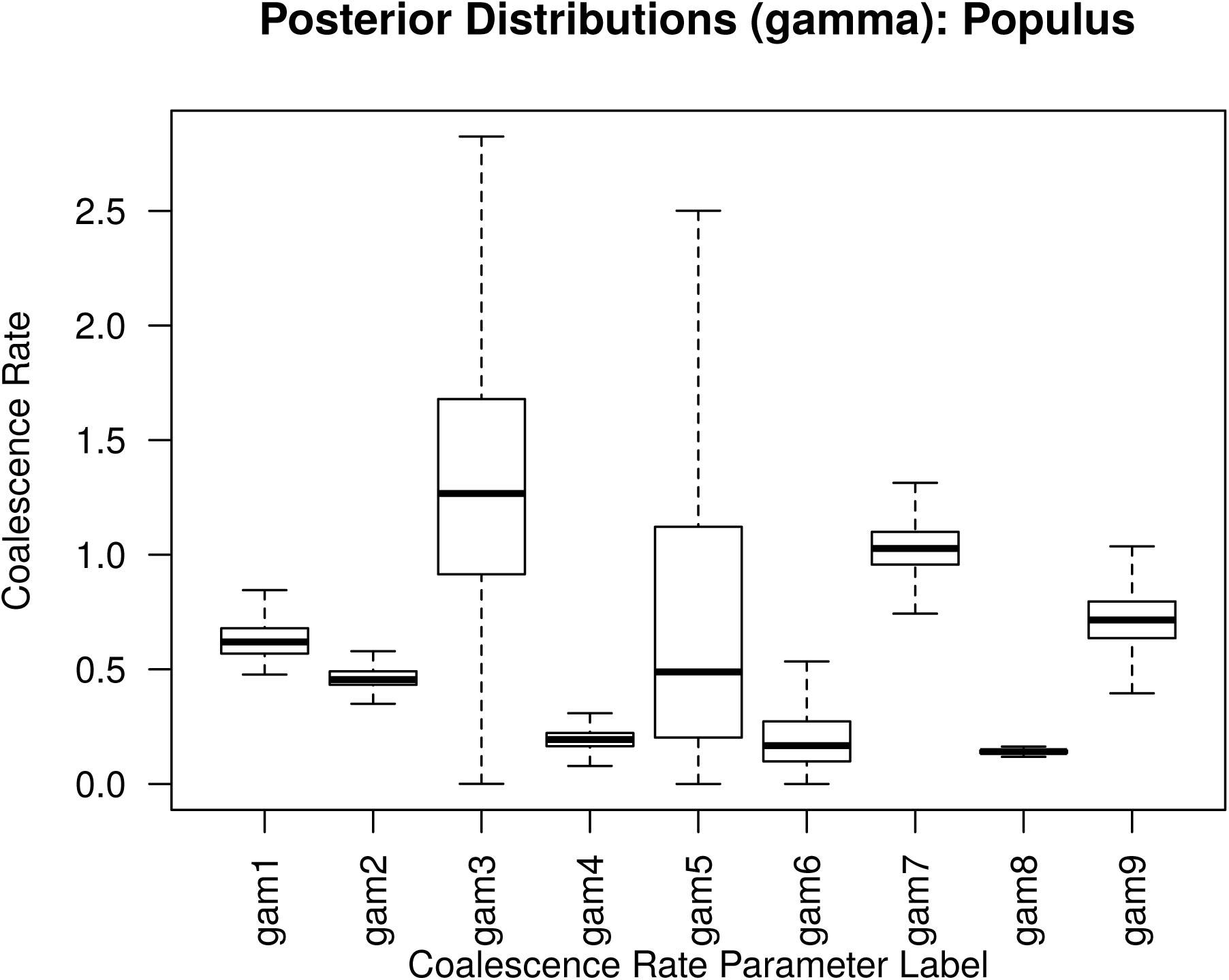
Posterior distributions of parameters for *Populus* data, using coalescence time inference with the model shown in Figure 8. Plot shows posterior medians (dark line), approximate middle 50% (box), and 90% (whiskers), for coalescence rates, labeled as in Figure 8. For instance, the coalescence rates for regions 4 and 8 (labeled “gam4” and “gam8”) are inferred to be much smaller than most other regions, perhaps indicating larger or older populations in southwestern British Columbia.

